# Isolated Grauer’s gorilla populations differ in diet and gut microbiome

**DOI:** 10.1101/2022.01.04.474987

**Authors:** Alice Michel, Riana Minocher, Peter-Philip Niehoff, Yuhong Li, Kevin Nota, Maya A. Gadhvi, Jiancheng Su, Neetha Iyer, Amy Porter, Urbain Ngobobo-As-Ibungu, Escobar Binyinyi, Radar Nishuli Pekeyake, Laura Parducci, Damien Caillaud, Katerina Guschanski

## Abstract

The animal gut microbiome has been implicated in a number of key biological processes, ranging from digestion to behavior, and has also been suggested to facilitate local adaptation. However, studies in wild animals rarely compare multiple populations that differ ecologically, which is the level at which local adaptation may occur. Further, few studies simultaneously characterize diet and the gut microbiome from the same sample, despite the likely presence of co-dependencies. Here, we investigate the interplay between diet and gut microbiome in three geographically isolated populations of the critically endangered Grauer’s gorilla, which we show to be genetically differentiated. We find population- and social group-specific dietary and gut microbial profiles and co-variation between diet and gut microbiome, despite the presence of core microbial taxa. There was no detectable effect of age, sex, or genetic relatedness on the microbiome. Diet differed considerably across populations, with the high-altitude population consuming a lower diversity of plants compared to low-altitude populations, consistent with food plant availability constraining diet. The observed pattern of covariation between diet and gut microbiome is likely a result of long-term social and ecological factors. Our study suggests that the gut microbiome is sufficiently plastic to support flexible food selection and hence contribute to local adaptation.

## Introduction

The ranges of many species span ecologically diverse habitats that differ in abiotic and biotic factors, leading to some degree of adaptation to the predominant local condition. Our view of how organisms adapt has recently expanded beyond natural selection acting on morphological, physiological, and behavioral traits, to also include the contribution of associated microorganisms, the microbiome (Rosenberg & Zilber-Rosenberg, 2016). In animals, the microbiome plays a critical role in key biological processes such as digestion, health, behavior (Agranyoni et al., 2021; Colston & Jackson, 2016; Davidson et al., 2020; Ley et al., 2008; Moran et al., 2019), and has even been implicated in influencing host genomic evolution (Rudman et al., 2019).

The gut microbiome is shaped by numerous factors including host evolutionary relationships, social interactions, habitat, and diet (Archie & Tung, 2015; Rojas et al., 2021; Youngblut et al., 2019). In wild animals, distinct populations living under different ecological conditions have frequently been shown to possess unique gut microbiomes (Bueno de Mesquita et al., 2021; Couch et al., 2020; Uren Webster et al., 2018). Along with spatial differences, studies often show shifts in the gut microbiome concordant with seasonal dietary changes (Baniel et al., 2021; Bergmann et al., 2015; Guo et al., 2021; Hicks et al., 2018). Such differences are expected, as microorganisms, with their large population sizes, rapid evolution, and flexible community structure, are able to react quickly to changes in environmental conditions (Koskella et al., 2017), supporting their role in local adaptation of the host (Alberdi et al., 2016). Experimental studies have used dietary and gut microbial manipulations to dissect the directionality of the diet-microbiome link. They suggest a two-way connection. On the one hand, dietary manipulations alter the composition of the gut microbiome, permitting hosts to rapidly utilize new dietary sources (Reese et al., 2021). On the other hand, the gut microbiome itself can drive dietary choice (Trevelline & Kohl, 2022). In the wild, it is possible that the microbiome may impact dietary choices by modulating host behavior, for example, by constraining the selection to similar foods even in different habitats or by promoting dispersal decisions that reduce environmental change (‘natal habitat-biased dispersal’).

Here, we investigate spatial variation of the gut microbiome and its potential role in local dietary adaptation by jointly analyzing dietary and gut microbial diversity and composition in several isolated populations of the critically endangered Grauer’s gorilla (*Gorilla beringei graueri*) (Plumptre et al., 2016). This gorilla subspecies is endemic to the eastern Democratic Republic of Congo (DRC). Our study populations occupy the ecological extremes of the species’ range, approximated here by altitude (600 m above sea level [asl] and 2500 m asl). Grauer’s gorillas are herbivores, consuming a large diversity of plants and plant parts (Yamagiwa et al., 2005). However, due to the political instability throughout their range, very little is known about ecology and diet of different populations (but see van der Hoek, Binyinyi, et al., 2021; van der Hoek, Pazo, et al., 2021).

Using fecal DNA metabarcoding combined with host genotyping, we first investigated whether isolated and genetically differentiated gorilla populations show dietary similarities. As plant communities differ considerably by altitude throughout the region (Imani et al., 2016), the presence of shared food taxa across populations would be indicative of restrictive dietary selection (a core Grauer’s gorilla diet). If such a pattern of food selection occurs at least in part via gut microbial influence over host foraging, we also expect to find a conserved set of gut microbial taxa (a core microbiome). In contrast, if plasticity of the gut microbiome confers dietary flexibility, potentially facilitating local adaptation, we expect diet and the microbiome to differ significantly among populations, with strong covariation between them and little evidence for conserved dietary and microbial taxa.

## Materials & Methods

### Ethics Statement

This study was conducted in compliance with legal requirements of the DRC and the animal use policies of UC Davis. Data collection protocols were approved by Institut Congolais pour la Conservation de la Nature. Samples were collected non-invasively, without disturbing the animals.

### Sample collection

Fecal samples (n=220) were opportunistically collected from Grauer’s gorillas in eastern DRC between 2015 and 2018 at three sites: Kahuzi-Biega National Park (KBNP, 2.32°S, 28.72°E; KBNP; 2500 m asl), Nkuba Conservation Area in Walikale territory, North Kivu (NCA, 1.38°S, 27.47°E; NCA; 600 m asl), and Maiko National Park (MNP, 0.87°S, 27.35°E; MNP; 830 m asl; **Figure 1**). In KBNP, gorillas in the Chimanuka group were habituated to human presence and samples were collected from identified individuals after observing defecation. All other samples were collected from night nests without knowledge of individual identity following the two-step collection method (Nsubuga et al., 2004). Geographic location and altitude were recorded using handheld GPS for all sampling sites except for the Mankoto group in KBNP, for which this information is missing. We assigned age classes in the field based on dung diameter, as follows: infant <4cm, sharing a nest with an adult; juvenile/subadult <5cm, own nest; and adults >5cm (McNeilage et al., 2006; Schaller, 1963). For the Chimanuka group, age classes of identified individuals were known from observations.

**Figure 1.**
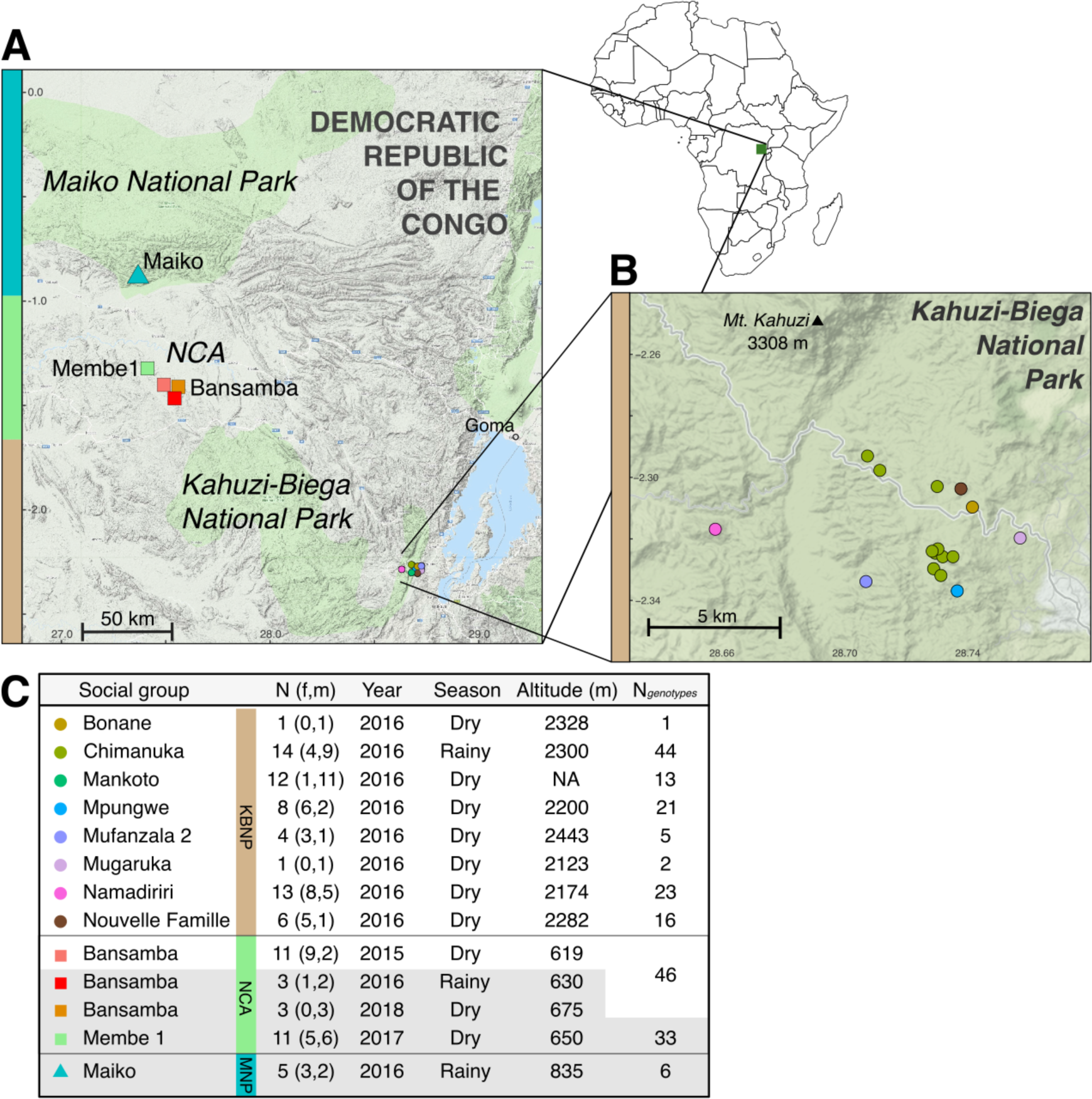
**(A)** Map of Grauer’s gorilla fecal sampling locations from Maiko National Park (MNP; designated with cyan on the left-hand side of the map), Nkuba Conservation Area (NCA; green) and Kahuzi-Biega National Park (KBNP; brown), with **(B)** inset zooming in on different social groups in KBNP. Circle colors designate social groups, coded as in **(C)**. Note that multiple circles are present for the Chimanuka group, consistent with opportunistic sampling of identified individuals. Geographic coordinates were not available for the Mankoto group. The table in **(C)** shows the sample size (N=number of unique individuals, f=number of females, m=number of males) used for dietary and gut microbiome characterization of each social group. Only diet but not gut microbiome data is available for samples shaded in gray. Bansamba group (NCA) was sampled repeatedly, but only few samples were included in dietary analyses in later years (three from 2016 and three from 2018). Also shown for each social group are: collection year and season, altitude, and N_genotypes_, the total number of successfully genotyped samples.

### DNA extraction

Fecal samples were exported to Uppsala University, Sweden, for molecular analysis. DNA was extracted from 50 mg of dried material using the DNeasy PowerSoil DNA Extraction Kit (Qiagen) in a dedicated primate fecal extraction laboratory. We implemented the following modifications to the manufacturer’s protocol: fecal samples were incubated under shaking (500 RPM) in the C1 solution overnight at 23°C. They were then transferred into a heating block and incubated at 65°C for 10 minutes, followed by bead beating on a vortex at maximum speed for 1 hour at room temperature. Incubation in C2 and C3 solution was on ice. We incubated the samples in C6 solution at room temperature for 5 minutes before elution.

### Gorilla genotyping, individual identification, relatedness and population differentiation analyses

We genotyped all 220 samples at 12 microsatellite loci (vWF, D1s550, D4s1627, D5s1457, D5s1470, D6s474, D6s1056, D7s817, D8s1106, D10s1432, D14s306, and D16s2624) following the two-step multiplex protocol (Arandjelovic et al., 2009) and sexed them with the amelogenin assay (Bradley et al., 2001). PCR products were run on an agarose gel to confirm amplification success and absence of contamination in blanks. Up to four loci were pooled, based on fluorophores and product sizes, and run on the ABI GeneAnalyzer (ThermoFisher Scientific). We scored genotypes manually in GeneMapper v5.0 (Chatterji & Pachter, 2006) and used Cervus v3.0.7 (Kalinowski et al., 2007) to identify individuals. Samples were considered to originate from the same individual if their genotypes matched at five or more loci without mismatches, with the probability of identity assuming full-sibling relationship (PIDsib) less than 0.05. We manually generated consensus individual genotypes from matching samples, taking into account the time and place of sample collection, and evidence about the presence of other individuals from the same group.

We tested for deviations from Hardy-Weinberg equilibrium, heterozygote deficiency, and linkage disequilibrium at each locus in GenePop v4.7.5 (Raymond & Rousset, 1995; Rousset, 2008). Genetic population structure was assessed using STRUCTURE v2.3.4 (Porras-Hurtado et al., 2013) with 20 independent runs for K = 1-11 (corresponding to the eleven social groups), an 100,000- iteration burn-in, and data collection for 1,000,000 runs, assuming population admixture and correlated allele frequencies (Falush et al., 2003). Results from different runs of K were merged in CLUMPP (Jakobsson & Rosenberg, 2007; Kopelman et al., 2015), and analyzed and visualized in ‘pophelper’ in R (Francis, 2017; R Core Team, 2021). The most likely value of K was determined using ΔK (Evanno et al., 2005). We used the ‘adegenet’ R package for Principle Component Analysis (PCA) based on individual genotypes (Jombart, 2008). Population differentiation statistics F_ST_ and F’_ST_ (Meirmans & Hedrick, 2011) were calculated in GenoDive v3.04 (Meirmans, 2020), and significance assessed with 9999 permutations. We compared genetic relatedness between populations and social groups using an AMOVA in the R package ‘poppr’ (Kamvar et al., 2014) and calculated pairwise relatedness (*r*) between all individuals within KBNP and NCA separately in ML-Relate (Kalinowski et al., 2006).

### Characterization of gorilla diet

We characterized the diet of 92 unique individuals identified by genotyping (see Results), from nine social groups and two lone silverbacks (**Table S1, S2**). We aimed to analyze a single nest site per group, but have also included individuals from additional nest sites of the same group if they were collected during the same year and season to maximize the number of studied individuals (**Table S2**). A single sample per individual was studied. The majority of our samples were collected during the dry season, but we also included some samples, social groups (Chimanuka) and one population (MNP) that were collected during the rainy season (**Table S2**). We present our analyses with and without these samples.

We amplified the P6 loop of the *trn*L chloroplast intron (Taberlet et al., 2007), a locus that has been successfully used for dietary metabarcoding in primates, and for which a large database of tropical plants is available (Mallott et al., 2018). We used the standard *trn*L g and h primers (**Table S3**), tagged with 96 eight-base-pair (bp) barcodes. Each barcode differed from all others at a minimum of three positions. DNA amplifications were carried out in 20 µl reactions containing 2 µl fecal DNA extract, 1 U Platinum II *Taq* Hot-Start DNA polymerase, 1x Platinum II Buffer, 0.2 mM each dNTP, 2 mM MgCl_2_, and 1 µM each primer. Each DNA sample was amplified twice. The duplicates were placed randomly on different PCR plates to avoid potential batch effects and biases due to cross-contamination of sample and/or barcoded primer (**Table S1**). We included one PCR negative and two to three empty wells per plate, to check for contamination during PCR (Taberlet et al., 2018). In addition, we included five DNA extraction blanks. PCR conditions consisted of 2 minutes denaturation at 94°C followed by 35 cycles of 94°C for 30 seconds, 51°C for 30 seconds, and 68°C for 15 seconds, without final extension. PCR products were checked on a 2% agarose gel to confirm amplification without contamination.

The barcoded PCR products were pooled column-wise (16 µl for each sample, duplicates in separate pools), mixed with 640 µl PB Buffer, and purified using MinElute columns (Qiagen, The Netherlands), eluting in 50 µl EB buffer. Double-indexed next-generation sequencing libraries (Kircher et al., 2012) were prepared as detailed (Brealey et al., 2020; Rohland et al., 2015) but using not-barcoded incomplete adapters after blunt-end repair. Two library preparation blanks were carried through all steps. Each pool was quantified using qPCR with PreHyb primers (**Table S3**; Rohland et al., 2015) and amplification settings as in Brealey et al. (2020).

Each sample pool and both library blanks received a unique combination of indices (**Table S1**). For indexing PCR, we used the same reaction mixture and cycling conditions as Brealey et al. (2020). The number of cycles ranged from 8 to 10, depending on the copy number estimated from qPCR (**Table S1**). Library preparation blanks were amplified for 10 cycles to maximize capture of potential contaminants. We performed MinElute purification and quantified indexed pools with qPCR, as above, using i7 and i5 primers (Rohland et al., 2015, **Table S3**). Indexed sample pools were combined in equimolar amounts, except for library preparation blanks, of which we added 0.5 µl each into the final pool, corresponding to the lowest amount added for any sample. The final sequencing pool was cleaned using AmPure XP beads (Beckman Coulter, USA) with two elutions (0.5x followed by 1.8x), which remove very large fragments and fragments <100bp, respectively. This size selection is optimized for the retention of *trn*L amplicons (∼10-150bp in length + 148 bp of barcoded and indexed adapters). Elution was performed in 30 µl of EB buffer. The cleaned library pool was quantified using both a Qubit High Sensitivity fluorometer and 2200 TapeStation and sequenced at the Uppsala Science for Life Laboratory on a single MiSeq lane with 150 bp paired-end sequencing with version 2 chemistry.

Sequence processing and analysis was done in OBITools v1.2.13 (Boyer et al., 2016). Paired reads with quality scores >40 and overlap >10 bp were retained and merged. Sample of origin for each read was established through its index and barcode, requiring an exact sequence match. Sequences were clustered into molecular operational taxonomic units (MOTUs), each representing a unique plant taxon (Valentini et al., 2009). A large number of MOTUs had fewer than 10 sequences across all samples and were removed as recommended (*e.g.*, Shehzad et al., 2012). We also removed sequences that differed by exactly one nucleotide from a more abundant sequence and had a total count less than 5% of the more abundant sequence, following Boyer et al. (2016).

Finally, taxonomic assignment used a custom-made reference database (below). Based on a frequency plot of identity to the reference database (**Figure S1**) and similar *trn*L-based studies of tropical primate diet (*e.g.,* Quéméré et al., 2013), we removed sequences below an identity threshold of 0.90. Below this, sequences were regarded as likely chimeric, enriched in sequencing or PCR errors. No singletons were present after this filtering step.

### Compiling plant trnL reference database

We built a local DNA barcoding reference library by downloading all 324,502 available sequences from NCBI GenBank using the search query: “(trnL[All Fields] OR complete genome[All Fields]) AND (plants[filter] AND (chloroplast[filter] OR plastid[filter]))” (last accessed 2 December 2021). In OBITools v1.2.13, the sequence list was annotated with taxonomy information downloaded from NCBI (ftp://ftp.ncbi.nih.gov/pub/taxonomy/taxdump.tar.gz, last accessed 3 December 2021). To complete the database of *trn*L genes, we followed established protocol (Boyer et al., 2016), using the same *trn*L g-h primers as in the wet laboratory to extract *trn*L variants *in silico* in the program ecoPCR v2.1 (Ficetola et al., 2010). We kept sequences that were between 10 to 230 base pairs long with at most three primer mismatches total (Taberlet et al., 2018). The final database contained 21,308 *trn*L *in silico* amplicons, in 608 families and 5,662 genera.

To evaluate the resolution of our reference database with respect to local plant diversity, we compared plant taxa present in our database to a list of plants known to occur in the Kahuzi and Itebero regions of KBNP (Yumoto et al., 1994). To enable this comparison, we updated the taxonomic classification of the KBNP plant list (Yumoto et al., 1994) by searching for species names in the Global Biodiversity Information Facility (GBIF). The updated list contained 328 taxa, in 81 unique families and 234 genera. Of these, all families and 77.4% of genera were present in our *trn*L database.

### Characterization of gorilla gut microbiome

We characterized gut microbial composition in 70 individuals from KBNP and NCA populations using a single sample per individual (**Table S2**). We selected the same sample that was used for dietary analyses and only dry season samples from the Bansamba group in NCA. To quantify possible contamination, we also carried nine random extraction blanks through the entire data generation process.

The V4 region of the *16S rRNA* gene was amplified with primers 515F/806R (**Table S3**) for each sample in duplicate. The PCR reaction contained 2 µl of extracted DNA, 5 µM each of the forward and reverse primer, 1x Phusion High-Fidelity Buffer, 0.02 units Phusion HF DNA polymerase (2U/µl), 0.012 mg DMSO and 0.05 µM (each) dNTPs, with the volume made up to 20 µl with Ultrapure H2O. Thermal cycling conditions were as follows: 30 seconds at 98°C, 25 cycles of 98°C for 10 seconds, 52°C for 20 seconds and 72°C for 20 seconds, and 10 minutes at 72°C. PCR cycles were limited to 25 to minimize the risk of unspecified products and chimeras. Duplicate reactions were pooled and cleaned with AmPure beads (Qiagen).

Next-generation sequencing libraries were prepared from PCR products following the double-barcoding, double-indexing strategy (Kircher et al., 2012; Meyer & Kircher, 2010; Rohland et al., 2015; van der Valk et al., 2017). As a result, each sample had a unique combination of two barcodes and two indices, which enabled bioinformatic filtering of potential chimeric molecules and misassigned reads resulting from index hopping (van der Valk et al., 2017, 2020). For indexing, we determined the suitable number of PCR cycles (8-11) based on qPCR of barcoded libraries, as above. Indexed libraries were quantified by qPCR and pooled in equimolar amounts for sequencing on a single MiSeq lane, using version 2 chemistry and 250 bp paired end sequencing at the Uppsala Science for Life Laboratory sequencing facility.

Sequencing reads were demultiplexed and adapters removed using a Python script (Brealey et al., 2021). We followed established protocol to estimate microbial amplicon sequence variants (ASVs) using DADA2 (Callahan et al., 2016), rather than clustering sequences, which avoids biases due to arbitrary similarity thresholds (Edgar, 2018). Forward and reverse reads were truncated to 200 and 150 bp, respectively, at which point read quality scores dropped below 35. We merged paired-end reads, requiring an overlap of at least 12 bp, and removed sequences outside the 250-256 bp range and those with any barcode mismatch, as recommended (Callahan et al., 2016).

Taxonomy was assigned using the SILVA 132 reference database, released in December 2017 (Quast et al., 2012). Species-level assignment required a strict 100% match (Edgar, 2018). We removed singletons and ASVs labeled ‘Unassigned’, ‘Eukaryota’, “mitochondria”, or “chloroplast”. We retained Archaea, although archaeal amplification from the V4 region of the *16S rRNA* is limited (Raymann et al., 2017), because within-dataset comparisons are nonetheless informative. We built a bacterial phylogenetic tree by aligning sequences to the Greengenes 13_5 mega-phylogeny (203,452 99% OTUs; DeSantis et al., 2006) in SEPP using default parameters (Mirarab et al., 2012).

### Statistical analyses of trnL and 16S datasets

To examine dietary and microbiome diversity, we analyzed the *trn*L and *16S rRNA* metabarcoding datasets, after first filtering out rare sequence variants below 0.5% relative abundance in at least one sample, as suggested (Deagle et al., 2019). We evaluated sampling effort and sequencing depth accumulation curves in the R packages ‘vegan’ (Oksanen et al., 2020) and ‘ranacapa’ (Kandlikar et al., 2018), respectively. We checked whether the predicted number of taxa (asymptote of the sequencing accumulation curve) minus actual number of taxa (richness) related to any of the considered biological variables or sequencing depth (read count) using a generalized linear model (GLM) with quasi-Poisson error distribution in the R package ‘lme4’ (Bates et al., 2015).

We calculated two alpha diversity metrics for each dataset: richness, or the number of taxa, and Shannon’s diversity index, or evenness (Chao et al., 2014). As recommended by McMurdie & Holmes (2013), we did not rarefy to minimum sequencing depth. To test the effects of population, social group, altitude, sex, and age class on diversity metrics, we fitted a GLM with quasi-Poisson (for richness) or gamma (for evenness) error distribution with logit link function, followed by Tukey honestly significant difference (HSD) *post-hoc* comparisons between levels of categorical variables that were overall significant (χ^2^ test with Bonferroni correction) (Lenth et al., 2021).

To assess trends in diet and microbiome beta diversity, or composition, we followed a strategy designed for the compositional nature of metabarcoding data (Gloor et al., 2017; Weiss et al., 2017). We used Bayesian multiplicative zero replacement and then centered and log-ratio (CLR) transformed each dataset using the R packages ‘zcompositions’ (Palarea-Albaladejo & Martín-Fernández, 2015) and ‘compositions’ (van den Boogaart & Tolosana-Delgado, 2008). For the microbiome dataset, we secondarily used Phylogenetic Isometric Log-Ratio Transform (phILR) to compute compositional abundance at phylogenetic balances (Silverman et al., 2017). To evaluate variation in composition of diet and microbiome, we computed Aitchison’s dissimilarity (Euclidean distance between CLR values) (Aitchison et al., 2000). To quantitatively estimate which factors best predict variation in diet and gut microbiome, we modeled the composition in CLR (or phILR) transformed space as a function of ecological and biological variables using PERMANOVA, via function *adonis2* in ‘vegan’ (Anderson & Walsh, 2013). The predictor variables were population, social group, sex, age class, and altitude. Sequencing read count was kept as the first predictor, even if p > 0.05. *Post-hoc* comparisons between levels of overall significant variables were done with Bonferroni correction using ‘pairwiseAdonis’ (Arbizu, 2020). The influence of genetic distance (*1 -* genetic relatedness) was modeled separately within each population using Mantel and partial Mantel tests (controlling for social group identity).

We estimated the covariance between diet and microbiome using a co-inertia analysis between the two matrices in the package ‘omicade4’ and calculated the RV coefficient (Escoufier, 1973; Robert & Escoufier, 1976) and its significance using a Monte Carlo test with 999 permutations (Meng et al., 2014). To compare the effects of diet and other variables on the gut microbiome, we fit a Multiple Regression on Matrices (MRM) model (Lichstein, 2007), an extension of the partial Mantel test, in ‘ecodist’ (Goslee & Urban, 2007). The explanatory variables were straight-line geographic distance, altitude difference, diet composition (Aitchison distance), and social group and population as binary (same, 0, or different, 1). Significance was assessed using 999 permutations of the response variable, the Aitchison distance matrix of gut microbiome composition.

Differences in beta diversity can be due to differential abundance of a few key organisms, or subtle differences across an entire community. To identify dietary and microbial taxa that may have driven compositional differences, we used the R package ‘ALDEx2’ and focused on significant differences (Wilcoxon rank sum test with correction for false discovery rate (FDR) *p* < 0.05) with effect sizes >1, as recommended (Gloor et al., 2017).

## Results

### Study populations of Grauer’s gorillas are genetically differentiated

We identified 92 unique individuals in the three study populations by microsatellite genotyping: 59 in KBNP, 28 in NCA, and 5 in MNP (**Figure 1, Table S2**). Individuals belonged to six different social groups and two solitary adult males (lone silverbacks) in KBNP, two social groups in NCA, and one group in MNP. Genotyping revealed that each individual was sampled 1-13 times, with 4-17 individuals per social group.

None of the 12 microsatellite loci deviated from Hardy-Weinberg equilibrium after Bonferroni correction for multiple testing (*p* > 0.1). On average, there were 6.1 alleles per locus (**Table S2**). The average observed and expected heterozygosities were 0.66 and 0.68, respectively. The test for global heterozygote deficiency was not significant overall (*p* = 0.6) or in any population (*p* > 0.4). The test for genotypic linkage disequilibrium using log likelihood ratio statistic with 66 pairwise comparisons between the 12 loci was not significant for any pair (*p* > 0.1). Thus, we assumed linkage equilibrium and considered all loci in further analyses.

Analysis of the three populations using STRUCTURE (Porras-Hurtado et al., 2013) revealed two distinct genetic groups (optimal K=2 according to ΔK; Evanno et al., 2005; **Figure S2**). The clusters differentiated gorillas in high-altitude KBNP (2500 m asl) from those in low-altitude NCA and MNP (600-830 m asl) (**Figure S3**), consistent with the PCA (**Figure 2A**). All three populations were significantly differentiated from one another (F’ST = 0.26-0.45; *p* < 0.001; **Table S4A**), with largest differences between MNP and KBNP, which are furthest apart geographically (215km). Individuals within social groups were more closely related than individuals in different groups in the same population (AMOVA ϕ = 0.12, *p* < 0.001; **Table S4B**), consistent with gorilla social structure (Harcourt & Stewart, 2013).

**Figure 2.**
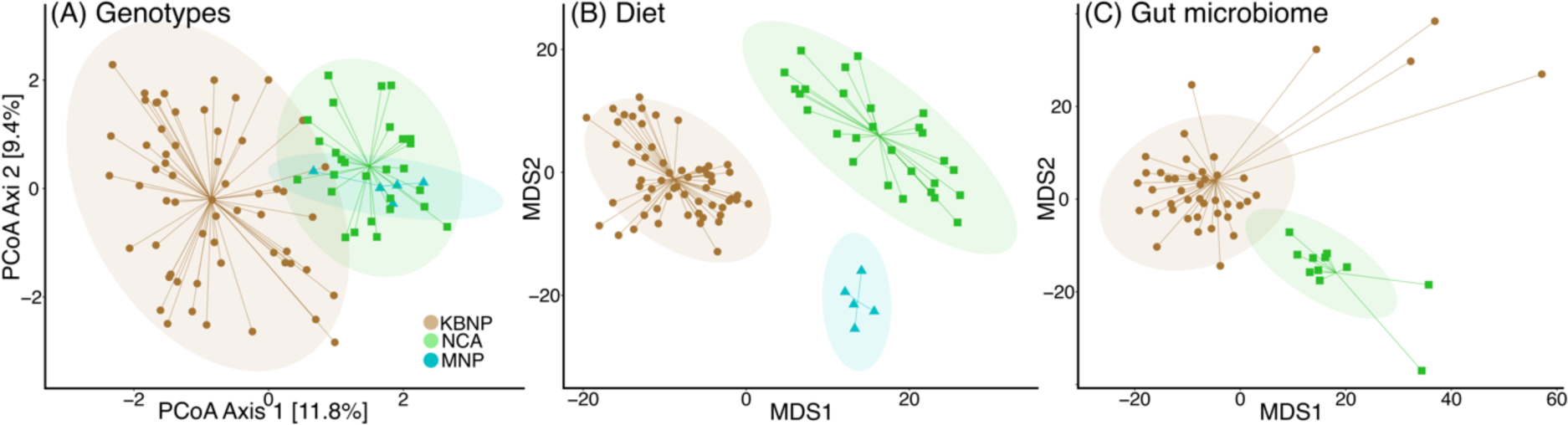
**(A)** PCA of genetic distances among individuals based on microsatellites. NMDS of **(B)** dietary composition and **(C)** gut microbiome composition, both in Aitchison distances. Individual samples are coloured by population of origin, with 95% confidence interval ellipses for each population (brown = KBNP, green = NCA, cyan = MNP, as in Figure 1).

### Negative controls in trnL and 16S rRNA metabarcoding

To quantify contamination in the diet (*trn*L) and the gut microbiome (*16S rRNA*) dataset, we analyzed DNA extraction blanks, PCR negative controls, unused barcode combinations and library preparation negative controls (for diet) (**Table S2**; **Table S5; Table S6**). In the diet dataset, the extraction and PCR negative controls contained 16 *trn*L reads in total, identified to 12 different plant taxa. Each taxon had one to three reads summed across all negative controls, yet up to 3,620-154,357 reads per sample (**Table S5**). There were no reads with unused barcode combinations, suggesting that cross-contamination of barcodes during PCR and library preparation was negligible. In the microbiome dataset, four extraction blanks had 90 reads in total, whereas the remaining five had none (**Table S6**). These mapped to eight different *16S* taxa, with three to 26 reads each. As with the diet data, these taxa were among the most abundant in the samples (up to 2,244-14,307 reads per sample). This pattern is consistent with low-level cross-contamination from high quantity into low quantity samples typical for large-scale sequencing studies (Eisenhofer et al., 2019).

### Diet of Grauer’s gorillas

We characterized the diet of 92 Grauer’s gorilla individuals (**Table 1C**) using the chloroplast *trn*L *P6 loop* locus. After data filtering, we retained 5,367,160 *trn*L sequencing reads (corresponding to 45% of raw reads) belonging to 120 unique taxa (**Table S7A**). PCR replicates were more similar to each other than to other samples in alpha and beta diversity (*p* < 0.001, **Figure S4**), and hence their sequencing data were pooled. Sample size and sequencing depth were sufficient to capture dietary diversity in KBNP and NCA, but not in MNP, where only five individuals were sampled (**Supplemental Text, Figure S5**).

**Table 1.** Top three most abundant plant taxa by population and taxa shared across populations.

| ID | NCBI- based finest taxonomic identity | Distribution-refined probable identity | Mean abundanc e in KBNP | Mean abund in NCA | Mean abund in MNP | KBNP Rank§ | NCA Rank§ | MNP Rank§ |
| --- | --- | --- | --- | --- | --- | --- | --- | --- |
| 1 | *Urera* sp. | *Urera hypselodendron* | 35.1% | 0.2% | 0.1% | 1 | 14 | 19 |
| 2 | Apocynaceae sp. | *Taccazea apiculata* | 21.0% | 0.2% | 0.2% | 2 | 16 | 17 |
| 6 | Urticaceae sp. | Urticaceae sp. | 6.0% | 12.9% | 0.06% | 3 | 13 | 22 |
| 8 | Myristicaceae sp. | *Pycnanthus*, *Staudtia*, or *Afradisia* sp. | 0.05% | 14.5% | 8.1% | 13 | 1 | 5 |
| 5† | Apocynoideae sp. | *Baissea*, *Funtumia*, or *Motandra* sp. | 4.6% | 8.9% | 14.7% | 4 | 2 | 3 |
| 25 | *Megaphrynium macrostachyum* | *Megaphrynium macrostachyum* | 0.01% | 3.4% | 0.05% | 25 | 3 | 26 |
| 31 | Phyllanthaceae sp. | Phyllanthaceae sp. | 0.01% | 0.01% | 18.8% | 46 | 63 | 1 |
| 32 | Alafinae sp. | *Strophanthus* sp. | 0.02% | 0.2% | 12.9% | 29 | 26 | 2 |
| 14† | *Ficus* sp. | *Ficus* sp. | 2.7% | 5.6% | 1.9% | 8 | 6 | 14 |
| 13‡ | Moraceae sp. | Moraceae sp. | 1.8% | 3.9% | 0.07% | 10 | 4 | 21 |

†Taxon greater than 1% relative abundance in all three populations (identifiable also by similar, high rank).
Shaded rows highlight taxa present in every sample collected during the dry season. With the exception of Moraceae sp. (ID 13‡), these taxa were also present in each rainy season sample and hence the abundances are shown including samples collected during the rainy season. Moraceae sp. (ID 13‡) was missing from one individual in the Chimamuka group (KBNP) collected during the rainy season.
§Rank is calculated by ranking each taxon by its relative abundance in a sample and calculating its mean rank across all samples in a population. It thus reflects the average abundance rank of a given taxon across all samples in a population.

Of the 120 detected dietary plant taxa, 115 could be identified to at least the order level (in 29 different orders), 110 to family (in 49 families), and 44 to genus (in 35 genera) level (**Table S8**). All but 21 taxa have previously been recorded in the Grauer’s gorilla diet in KBNP, NCA, and Mt. Tshiaberimu (Kambale, 2018; van der Hoek, Pazo, et al., 2021; Yamagiwa et al., 1994, 2005; Yumoto et al., 1994; **Table S8, columns S-T; Figure S6**). These 21 taxa are, however, present in the region (Spira et al., 2018).

Each Grauer’s gorilla fecal sample contained 36 – 80 *trn*L taxa (mean 58.52 ± 10.83) (**Figure 3A**), with each population showing a different set of most abundant and prevalent plants (**Table 1**; **Table S8**). Five plant taxa were found in each sample collected during the dry season in KBNP and NCA, even though they showed very low abundance in some samples (0.1%). Only two plant taxa had abundances over 1% in all three populations (**Table 1**).

**Figure 3.**
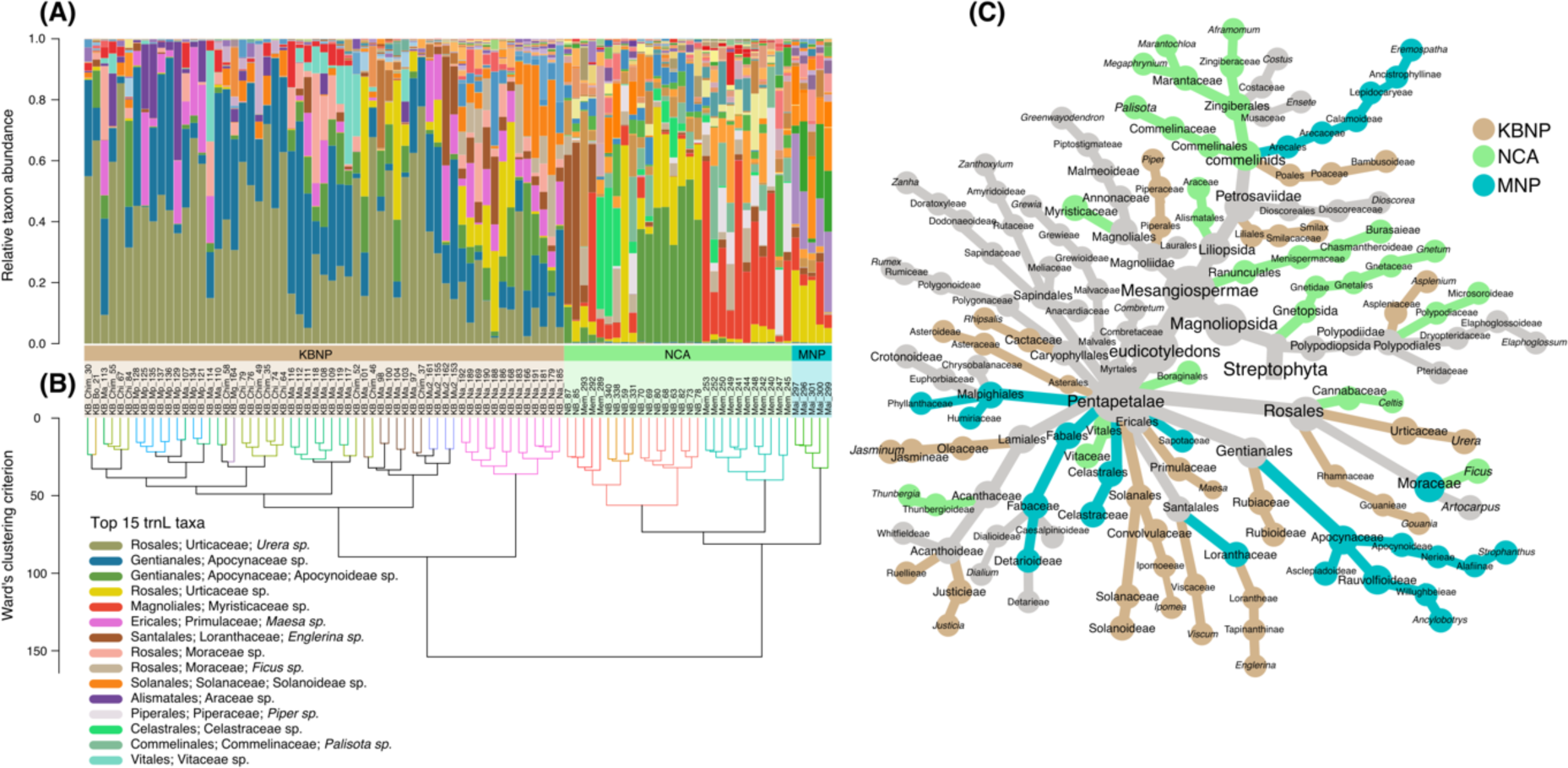
**(A)** Plants consumed by Grauer’s gorillas in KBNP, NCA, and MNP. The 15 most abundant taxa across all samples are shown. Populations are designated with coloured bars below (MNP cyan, NCA green, KBNP brown). **(B)** Hierarchical cluster dendrogram of Ward’s sum of squares based on minimum variance of squared dissimilarities (Murtagh & Legendre, 2014) of centered-log-ratio (CLR) transformed taxon abundance. Branches are colored by social group, following the code in Figure 1. **(C)** Plant taxa in Grauer’s gorilla diet, coloured by the population in which they are significantly more abundant (ALDEx2 Wilcoxon test p < 0.05). For taxa that differ between two or more population pairs, the color corresponds to the population with greatest effect size. Gray taxa do not differ significantly in abundance between populations. Branch lengths do not reflect phylogenetic distance. Diagram generated with the ‘metacoder’ package in R (Foster et al., 2017).

### Geography, altitude, and social group identity influence dietary diversity and composition in Grauer’s gorillas

Dietary richness and evenness differed significantly by population and social group identity (*p* < 0.001, **Table S9**). Both richness and evenness were significantly higher in low altitude populations (NCA and MNP) than in high-altitude KBNP (mean richness: 66.8±7.5 taxa in MNP, 65.6±6.1 in NCA vs. 54.5±10.4 in KBNP, *p* < 0.001; evenness: 10.0±1.3 in MNP, 8.4±2.5 in NCA, vs. 5.4±2.6 in KBNP, *p* < 0.001; **Figure S7**). Altitude was also a significant predictor of dietary richness and evenness in KBNP (p < 0.001; **Figure S8**). In contrast, neither individual’s sex nor age (age class available for 70 individuals) had an effect on dietary richness or evenness (*p* > 0.3; **Table S9**). We obtained qualitatively similar results when analyzing only samples collected during the dry season (excluding Chimanuka group, three individuals from Bansamba group, and MNP; **Table S9**), with the exception that dietary richness did not significantly change with altitude in KBNP (*p* = 0.2).

Hierarchical clustering of dietary composition first separated high altitude (KBNP) from low altitude (NCA and MNP) locations (**Figure 3B**), even though MNP samples were collected during the rainy season. Within populations, individuals clustered by social group. NMDS ordination showed a similar pattern (**Figure 2B**). After accounting for sequencing depth, dietary composition was significantly influenced by population (*p* < 0.001, explaining 27.9% of the variance) and social group (*p* < 0.001, explaining an additional 21.6%) but not by sex (*p* = 0.7) or age (*p* = 0.2; **Table 2**). All social groups differed significantly from each other (*p* < 0.05; **Table 2**), except for some comparisons involving the Mufanzala2 group, which had a small sample size (n=4). Restricting the analysis to two similarly sized social groups in NCA and KBNP collected during the dry season, we confirmed the presence of significant between-group and between-population diet differences (**Table S10**), supporting the notion that populations and social groups have distinct diets and that our results are not driven by differences in sample size or season.

**Table 2.**
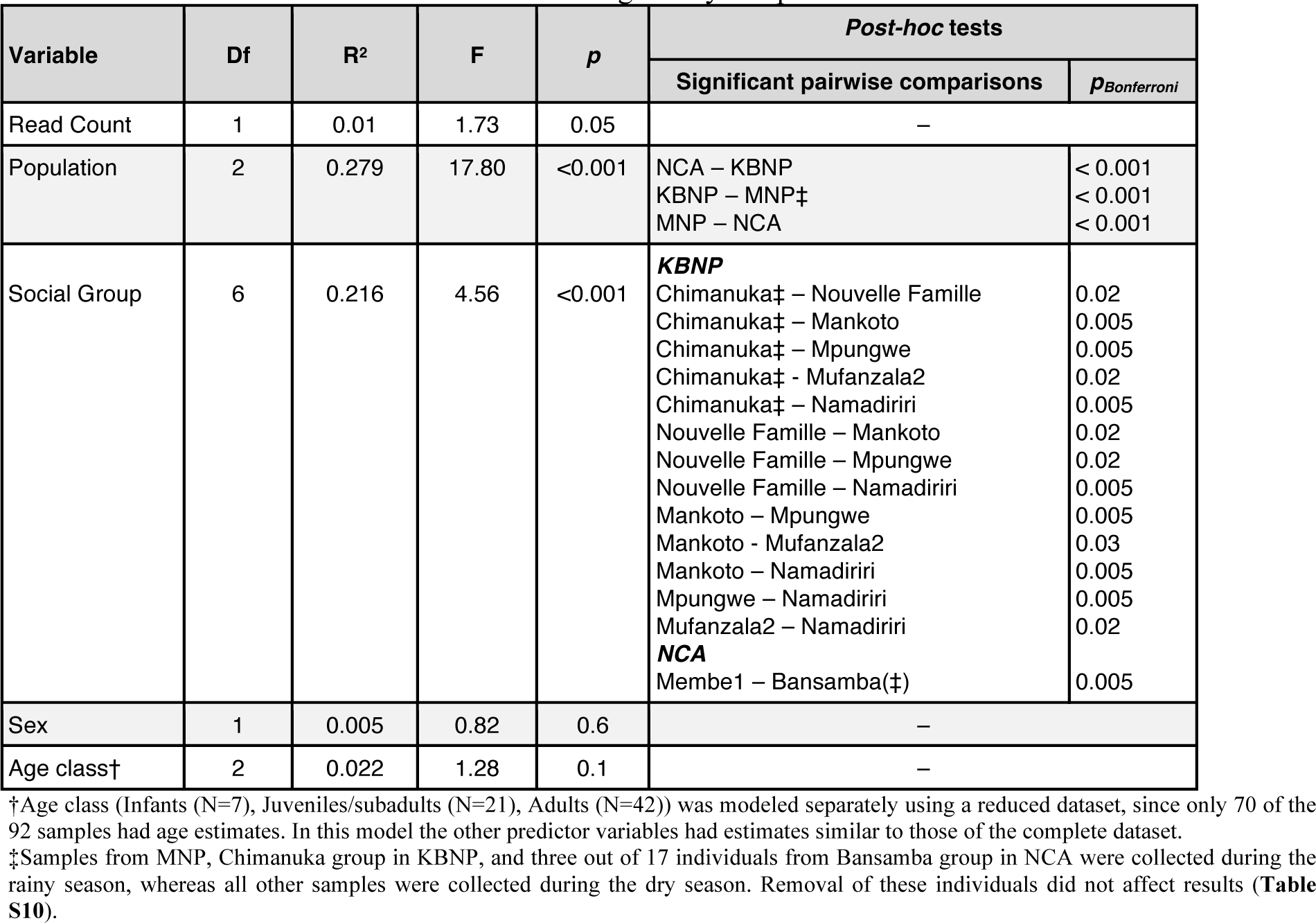
PERMANOVA model of factors influencing dietary composition

Using ALDEx2, we identified differentially abundant dietary items across populations. The drivers of the observed population dietary differences were among the most abundant taxa in each population (**Table 1**), most of which were absent or present at very low abundance in other populations (**Figure 3C**; **Table S11**). Out of the 21 previously undescribed food items (see above), 13 were significantly more abundant in low-altitude populations compared to the high-altitude population KBNP (**Table S8, S9; Figure S6**). Within populations, each social group consumed between two and 32 differentially abundant taxa (mean = 11.3 ± 11.6).

### Gut microbiome of Grauer’s gorillas in Kahuzi-Biega National Park and Nkuba Conservation Area

We characterized *16S rRNA* diversity in 70 individuals for which we also had dietary data (**Figure 1C**; **Table S2**), using the same samples as for diet. Two samples had low read counts (5 and 348, compared to the mean of 43,611 ± 11,357 in other samples) and were excluded. Our final dataset consisted of 68 unique individuals and contained 2,965,516 reads in 417 unique microbial taxa (**Table S12**).

The sample accumulation analyses suggested that additional sampling of feces from more individuals could uncover novel gut commensals at the population level (**Figure S9A**). However, per sample sequencing depth was sufficient to obtain a good representation of host microbiome diversity (**Figure S9B**). We detected 16 different phyla and 48 different families of microorganisms in the gut microbiome of Grauer’s gorillas. All taxa were identified at least to the family level, 309 taxa to the genus and 17 to the species level (**Table S12**). None were closely related to dominant soil microorganisms (Delgado-Baquerizo et al., 2018). There were seven Archaea in our dataset, belonging to the Methanomethylophilaceae and Methanobacteriaceae families. Each fecal sample contained on average 200.29 ± 19.6 taxa (min = 160, max = 237), each with average abundance of 0.2% ± 0.4%. Eleven taxa were present in every individual gorilla fecal sample from both populations (the core gut microbiome), however, populations differed in the most abundant taxa (**Table 3**).

**Table 3.**
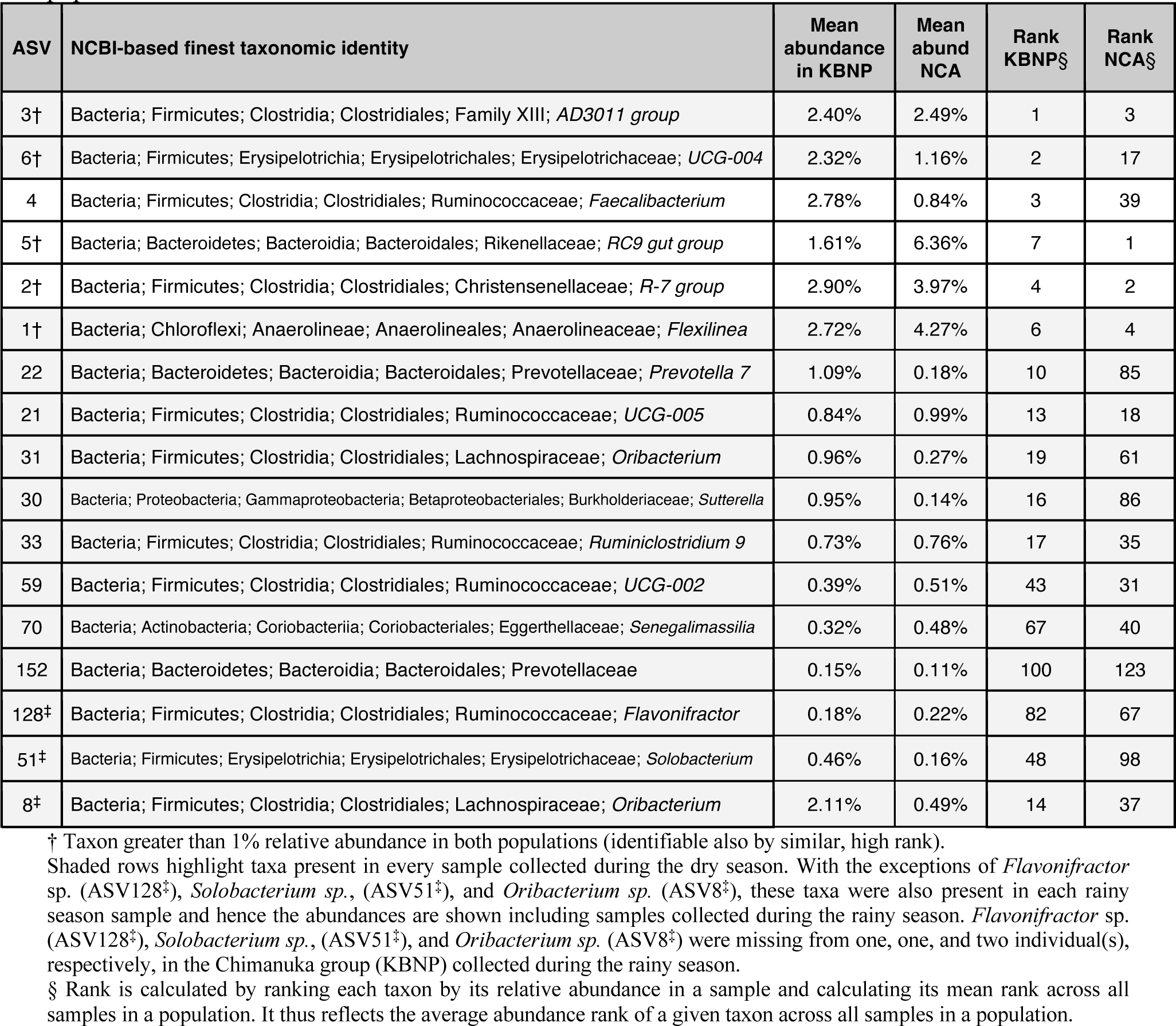
Top three most abundant gut microbiome taxa by population and taxa shared across populations.

In accordance with previous studies on great apes (Campbell et al., 2020; Gomez et al., 2016b; Hicks et al., 2018; Nishida & Ochman, 2019), Grauer’s gorilla gut microbiome in both populations was dominated by the phyla Firmicutes (65.6% in KBNP, 60.0% in NCA%), Bacteroidetes (20.7% in KBNP, 23.1% in NCA.0%), Spirochaetes (3.5% in KBNP, 5.4% in NCA), Chloroflexi (2.7% in KBNP, 4.3% in NCA), Proteobacteria (2.8% in KBNP, 3.4% in NCA), and Actinobacteria (2.0% in KBNP, 1.8% in NCA) (**Figure S10**), representing a diversity of microbial families (**Figure 4A**).

**Figure 4.**
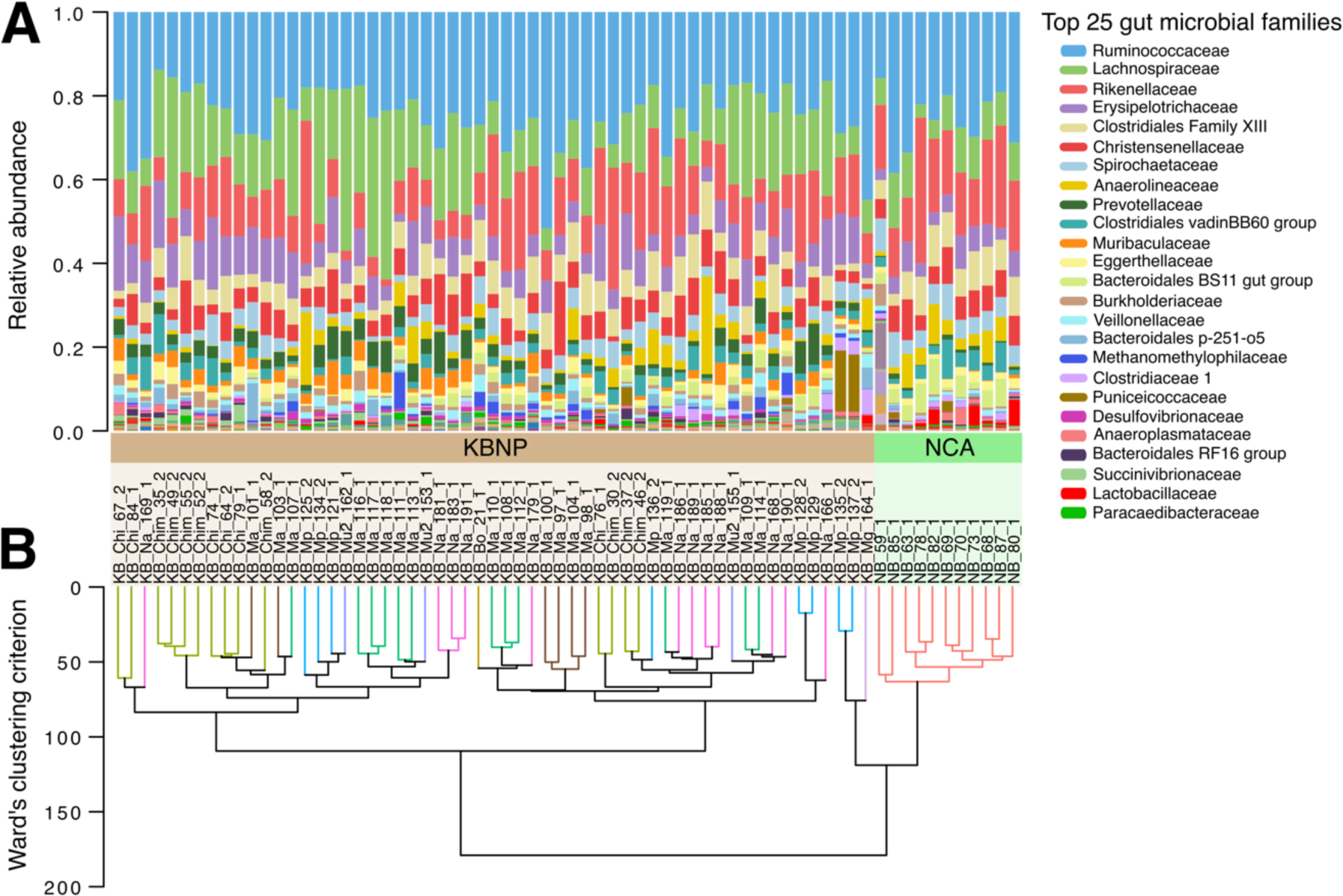
Gut microbiome composition **(A)** at the family level and **(B)** showing population clustering in composition, using CLR Aitchison distances dendrogram based on Ward’s clustering criterion (Murtagh and Legendre 2014).

### Diversity and composition of the gut microbiome in Grauer’s gorillas correlates with population and social group identity

Gut microbiome richness and evenness were significantly higher in the high-altitude population (richness: mean KBNP = 202.2±20.0 taxa vs. NCA = 190.2±14.0; *p* = 0.02; evenness: 83.7±17.8 vs. 74.2±12.8, *p* = 0.01), the opposite trend to diet, although neither microbiome richness nor evenness were related to altitude within KBNP (*p* = 0.07, 0.9; **Table S13**; **Figure S11**). While richness of the microbiome did not differ by sex (*p* = 0.2), females had more even microbiomes than males (85.7 vs. 78.5, *p* = 0.002). There were no differences by age (richness *p* = 0.3; evenness *p* = 0.1). The gut microbiome alpha diversity differed significantly by population even after removing rainy season samples (excluding Chimanuka group; richness *p* = 0.001, evenness *p* = 0.008; **Table S13**).

Gut microbiome composition differed between the two populations (KBNP and NCA) (**Figure 2C**) and among social groups (**Figure 4A-B**), with population explaining 10.5% of the total variance, and social group in KBNP explaining an additional 17.8% (*p* < 0.001; **Table 4**). Intergroup differences were significant, including among groups collected during the dry season (**Table 4**). Overall, gut microbiome dissimilarity was largest between individuals of different populations, followed by individuals from different social groups, and smallest between individuals from the same social group (**Figure 5C**). While altitude explained 12.7% of the variance across populations (N=56, *p* < 0.001) it only accounted for 3.8% in KBNP (N=45, *p* = 0.01). Genetic distance among gorillas was not a significant predictor of gut microbiome composition in NCA (N=11, ρ = -0.080, *p* = 0.7) or KBNP when social group was also considered (ρ = 0.015, *p* = 0.3). Microbiome composition did not differ by sex or age (*p* > 0.05, **Table 4**). Results using only dry season samples (**Table S14A**) and phylogeny-informed (phILR) distances were qualitatively similar (**Table S15**).

**Figure 5.**
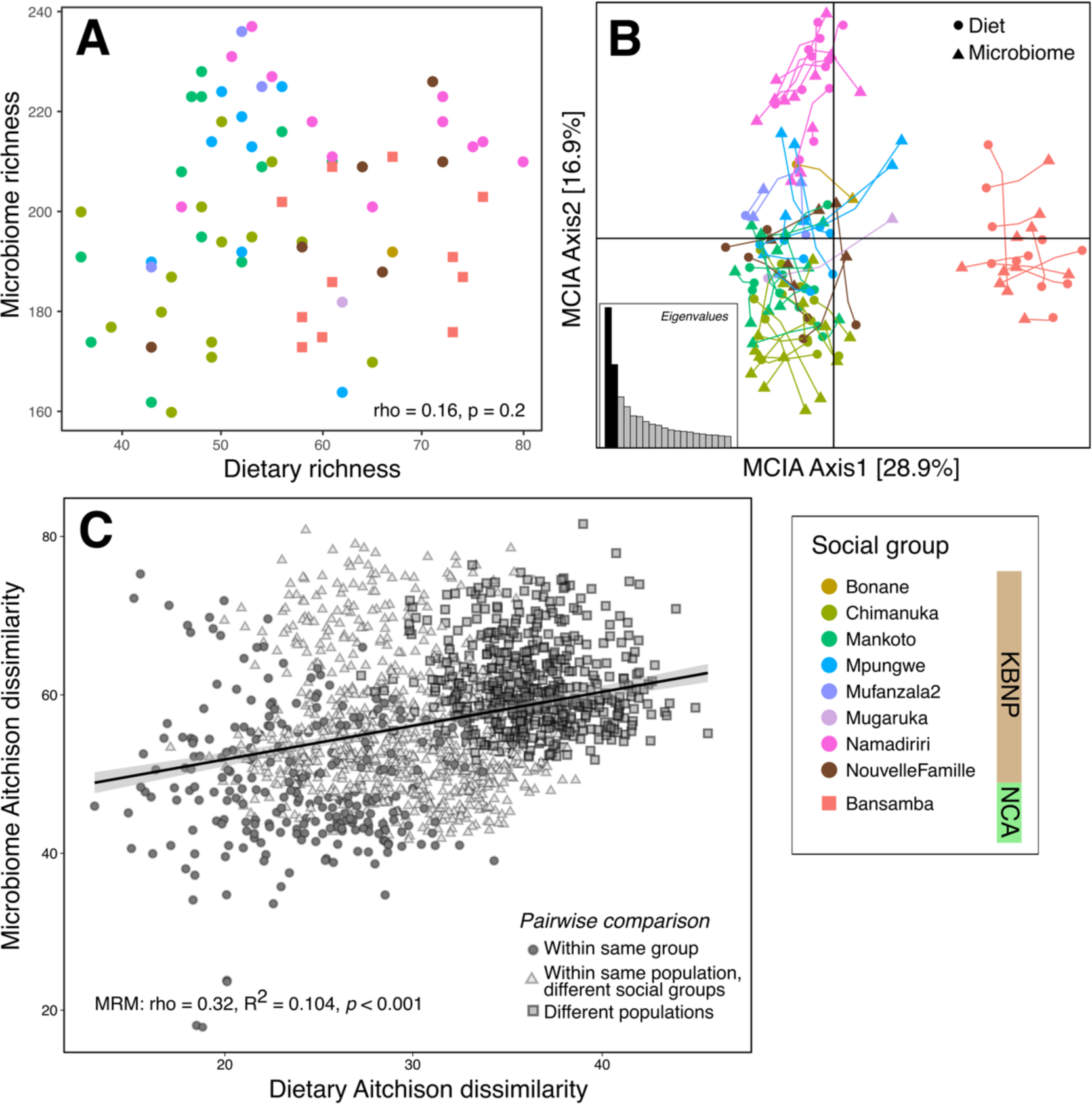
Relationship between diet and gut microbiome. **(A)** Microbiome and dietary richness, assessed as per-sample total sequence count, are not correlated (p = 0.2). **(B)** High multiple co-inertia (MCIA) between microbiome and diet composition in CLR-transformed space with Aitchison distance (RV = 55.7%, MC *p* < 0.001 based on 999 permutations). **(C)** Compositional differences (Aitchison distances) in diet and microbiome between samples (i.e., individual gorillas) are correlated in matrix regression.

**Table 4.**
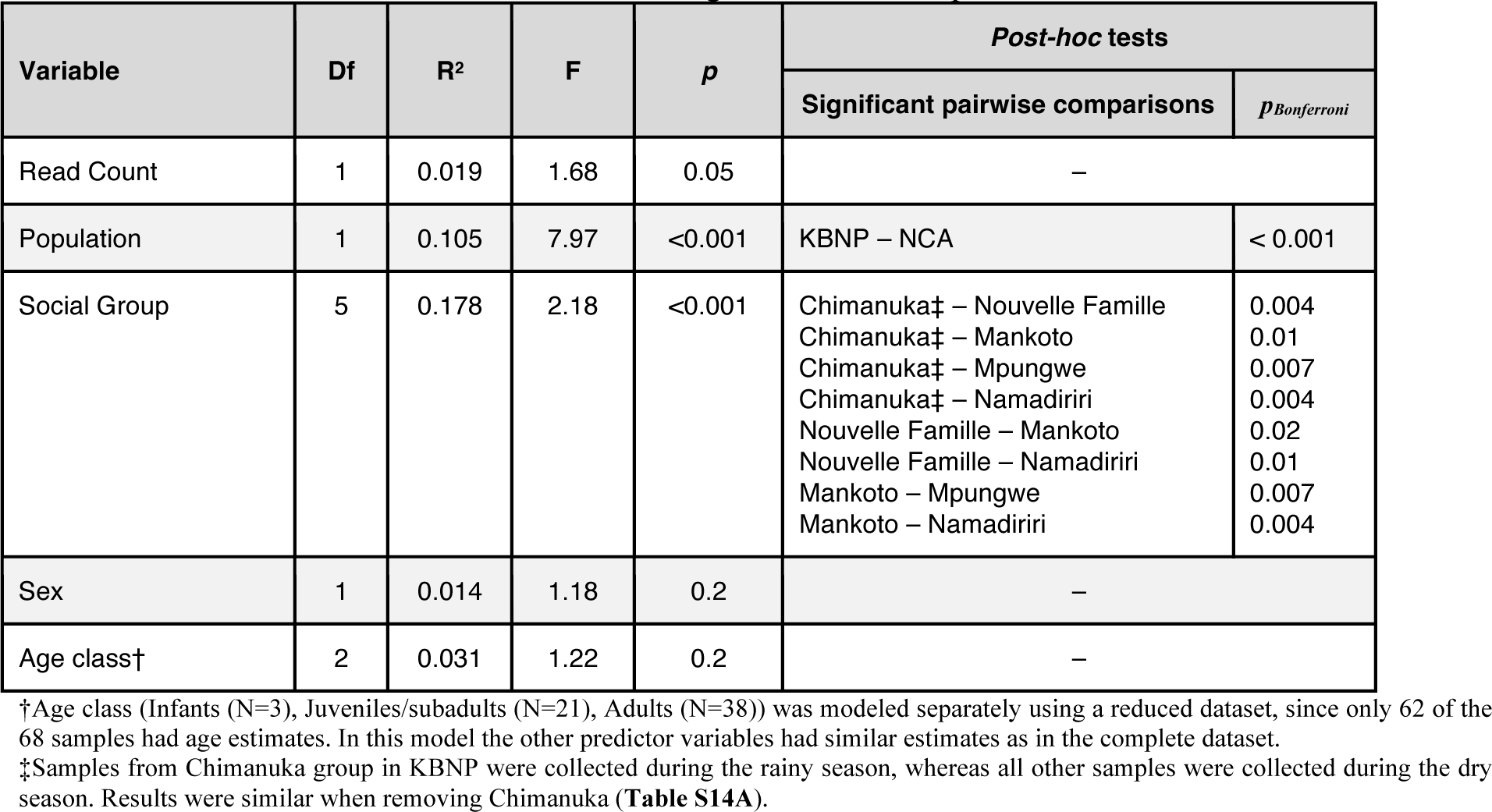
PERMANOVA model of factors influencing microbiome composition

| Variable | Df | R<sup>2</sup> | F | p | Post-hoc tests |  |
| --- | --- | --- | --- | --- | --- | --- |
|  |  |  |  |  | Significant pairwise comparisons | p<sub>Bonferroni</sub> |
| Read Count | 1 | 0.019 | 1.68 | 0.05 | – |  |
| Population | 1 | 0.105 | 7.97 | <0.001 | KBNP – NCA | < 0.001 |
| Social Group | 5 | 0.178 | 2.18 | <0.001 | Chimanuka‡ – Nouvelle Famille<br>Chimanuka‡ – Mankoto<br>Chimanuka‡ – Mpungwe<br>Chimanuka‡ – Namadiriri<br>Nouvelle Famille – Mankoto<br>Nouvelle Famille – Namadiriri<br>Mankoto – Mpungwe<br>Mankoto – Namadiriri | 0.004<br>0.01<br>0.007<br>0.004<br>0.02<br>0.01<br>0.007<br>0.004 |
| Sex | 1 | 0.014 | 1.18 | 0.2 | – |  |
| Age class† | 2 | 0.031 | 1.22 | 0.2 | – |  |

†Age class (Infants (N=3), Juveniles/subadults (N=21), Adults (N=38)) was modeled separately using a reduced dataset, since only 62 of the 68 samples had age estimates. In this model the other predictor variables had similar estimates as in the complete dataset.
‡Samples from Chimanuka group in KBNP were collected during the rainy season, whereas all other samples were collected during the dry season. Results were similar when removing Chimanuka (**Table S14A**).

We identified 42 taxa that significantly differed in abundance between NCA and KBNP (*p* < 0.05, effect size > 1) (**Table S16**). At the family-level, gorilla gut microbiomes in KBNP had a higher abundance of Muribaculaceae and Erysipelotrichaceae, whereas the gut microbiomes in NCA had more Spirochaetaceae and Christensenellaceae. At a finer phylogenetic level, populations differed in abundance of specific ASVs belonging to common, shared families like Rikenellaceae, Lachnospiraceae, and Ruminococcaceae.

### Diet and gut microbiome co-vary across studied populations

Compositional differences in dietary and gut microbial profiles showed significant co-inertia (RV = 0.557, *p* < 0.001; **Figure 5B**) and were correlated (ρ = 0.32, *p* < 0.001; **Figure 5C**), even after removing the rainy season samples from Chimanuka (RV = 0.599, p < 0.001; ρ = 0.35, *p* < 0.001). We detected no correspondence between dietary and gut microbial richness (*p* = 0.2; **Figure 5A**) or evenness (*p* = 0.1). Microbiome composition is known to change with diet in individuals and also differs between populations with different dietary preferences (*e.g.,* Reese et al., 2021; Youngblut et al., 2019). However, in our dataset, only population and social group were significantly correlated with gut microbiome composition, whereas dietary composition, geography, and genetic relatedness had no effect after accounting for social group and population of origin (**Table 5; Table S14B**). As with other analyses, dataset subsampling indicated that results were robust to sample size differences between populations (**Table S17**).

**Table 5.** MRM model comparing the effects of geography, diet, and sociodemographic factors on Grauer’s gorilla gut microbiome composition^‡^

| MRM Model of Gut Microbiome Composition |  |  | MRM Model Statistics |  |  |
| --- | --- | --- | --- | --- | --- |
| Explanatory Variable | Spearman's $\rho$ | $p$ | N | R<sup>2</sup> | F-statistic |
| Geographic distance | 0.07 | 0.4 | 56 | 0.278 | 98.35 |
| Altitudinal difference | -0.08 | 0.6 |  |  |  |
| Diet composition† | -0.19 | 0.07 |  |  |  |
| Population | 0.64 | < 0.001 |  |  |  |
| Social group | 0.45 | < 0.001 |  |  |  |

† Microbiome and diet composition in Aitchison distances.
‡ Model results without Chimanku are shown in Table S14B.

## Discussion

In this study, we applied fecal genotyping and DNA metabarcoding to identify Grauer’s gorilla individuals and characterize their diet and gut microbiome in three populations from across the species’ range. We were able to include a so far unstudied population from MNP and show that it is genetically distinct from two previously assessed populations KBNP and NCA (Baas et al., 2018). Grauer’s gorillas occur across the greatest altitudinal range of all gorilla taxa (Plumptre et al., 2016) and our study sites include the low and high altitude extremes. This provided us with the opportunity to test for dietary and gut microbial co-differentiation among the isolated populations of this critically endangered great ape. In particular, we set out to investigate if the gut microbiome may facilitate local adaptation by supporting digestion of diverse foods. Alternatively, the presence of conserved dietary patterns across populations along with a core gut microbiome would be indicative of a stabilizing role of gut microorganisms, which may limit ecological adaptation. Gorillas consume a wide variety of herbaceous vegetation, and fruits, when available, and their diet shows seasonal variation (Harcourt & Stewart, 2013; Rogers et al., 2004; Rothman et al., 2008; Yamagiwa et al., 1994). In Grauer’s gorillas, studies that rely on different methodologies suggested some differences in diet between populations (van der Hoek, Pazo, et al., 2021; Yamagiwa et al., 2005). However, previous studies did not assess the gut microbiome, and hence could not characterize its contribution to these differences.

Our joint diet and gut microbiome analyses provide little evidence for dietary conservation across populations but uncover a stable set of gut microorganisms that are shared among geographically, genetically, and ecologically distinct populations of Grauer’s gorillas. We detect co-variation in diet and microbiome, likely as a result of habitat differences and social factors among populations and social groups. Our results are thus consistent with the notion that the gut microbiome, although being conserved to some degree, provides sufficient flexibility to allow exploitation of diverse dietary resources and hence could contribute to local adaptation. In addition, we obtain evidence that dietary choice in Grauer’s gorillas is at least partially determined by plant availability, with a larger dietary repertoire at lower elevations.

### New insights into Grauer’s gorilla diet and feeding behavior

Grauer’s gorillas in the three study populations consumed 120 different plant taxa (**Table S8**), which is similar to the dietary composition and diversity reported in observational studies (116 and 100 different plants; van der Hoek, Pazo, et al., 2021; Yamagiwa et al., 2005; respectively). Low altitude populations consumed a greater diversity of plants than high altitude populations (**Figure 3**; **Figure S7**; **Table S9**), consistent with higher biodiversity (including plant diversity) at lower altitudes (Imani et al., 2016; Rahbek, 1995). We documented an average of 54-66 different plant taxa in each fecal sample, which is considerably more than reported daily diversity of consumed plants per individual based on behavioral observations (17 plant taxa per day on average in KBNP; Yamagiwa et al., 2005). In captivity, gorilla gut retention time was estimated to be 24 to 60 hours (Remis, 2000). Therefore, each fecal sample could represent plants consumed over the period of up to three days, with some items digested faster than others. Alternatively, our method could capture taxa that are missed in observational studies because they are consumed infrequently or in small quantities, at times of the day that are rarely observed (early in the morning or late in the evening), or which may be contaminants, parts of nest building material or involved in play or display and unrelated to diet. We discuss other potential limitations of molecular dietary analyses in detail below.

We detected 21 plant taxa that have, to our knowledge, not been reported as Grauer’s gorilla foods (**Table S8**; **Figure S6**). Some of these plants grow in KBNP (Spira et al., 2018) and are consumed by mountain gorillas (*e.g.,* Solanoideae; Rothman et al., 2014; Watts, 1984) or western lowland gorillas (*e.g.,* Laruales; Remis et al., 2001). Other plants, such as *Gnetum* and Humiriaceae, have not been documented in KBNP but are western lowland gorilla foods (Rogers et al., 2004; Takenoshita & Yamagiwa, 2008), which is consistent with their significantly higher abundance in the low-altitude sites of MNP and NCA.

Grauer’s gorillas in different populations consumed distinct diets (**Figure 2, 3**; **Table 1, 2**), with only two taxa shared across all three populations at an average abundance >1% per sample: *Ficus sp*. and Apocynoideae sp. (likely *Baissea sp., Funtumia sp.*, or *Motandra sp.* based on plant distribution; Spira et al., 2018). At broader taxonomic scales, all individuals consumed four plant families (Urticaceae, Apocynaceae, Moraceae, and Vitaceae), but the relative abundances varied considerably across populations, from less than 1% to up to 42%. The detection of shared taxa suggests that the same plants or their close relatives are present in the habitat of all three populations. However, the pronounced differences in their relative abundance suggest either that (1) their availability differs across study sites, and gorilla dietary choice is essentially passive and primarily based on food availability, or (2) that gorilla dietary choice is strongly determined by social factors, and food selection is a result of variation in culturally-transmitted feeding preference that differ across populations and social groups. Higher dietary diversity of low-altitude populations supports the first notion of rather opportunistic consumption of available plants. However, we also uncover distinct dietary signatures of social groups from the same population, which is consistent with social factors playing a role. Since gorilla groups show extensive range overlap, they would be well-suited for future investigations into the role of cultural versus ecological factors affecting dietary choices by evaluating if group-specific dietary patterns persist even when different social groups use the same habitat.

### The role of Grauer’s gorilla gut microbiome in ecological adaptation

In accordance with previous studies (Amato et al., 2019; Campbell et al., 2020; Gomez et al., 2016b; Moeller et al., 2014), we detect evidence for the presence of a Grauer’s gorilla core gut microbiome (**Table 3**, **Figure 4**). We identified eleven taxa belonging to cellulose- and other carbohydrate-degrading clades that were present in all study samples. Many of the microbial phyla, families, genera that are conserved across Grauer’s gorilla samples are also common in other great ape gut microbiomes, including western lowland gorillas, chimpanzees, and humans (Campbell et al., 2020; Fontsere et al., 2021; Gomez et al., 2015, 2016b; Hicks et al., 2018; Nishida & Ochman, 2019). Despite shared taxa, the gut microbiome composition in Grauer’s gorillas differed by population and to a lesser extent by social group, but considerably less so than dietary composition. This could be explained by the functional constraints placed on the gut microbiome, with key taxa required to perform essential functions in digestion. Other taxa may be allowed to vary and co-diversify with the host. Indeed, inter-species studies in primates find a strong effect of host evolutionary relationships on gut microbiome structure and composition (Amato et al., 2019) and we expect to detect similar, albeit less pronounced differences across isolated populations.

Our results indicate that dietary choice is not constrained by the gut microbiome. This is in line with many studies showing that animals, including all subspecies of gorillas (Harcourt & Stewart, 2013), experience seasonal dietary changes, which are also accompanied by gut microbiome changes (Baniel et al., 2021; Gomez et al., 2016a; Hicks et al., 2018; Orkin et al., 2019; Sharma et al., 2020). Here we detect differences between isolated populations, sampled during the same season, which are likely the joint result of gut microbiome-host co-diversification and plasticity of the gut microbial community that may facilitate local adaptation to different environmental conditions. Inter-population differences tended to derive from differential abundance of specific taxa within common bacterial families, like Lachnospiraceae and Rikellenaceae, which is consistent with the presence of the core microbiome in our study populations. There were, however, several differences at the family level that exemplify microbiome plasticity. For example, compared to KBNP, *Treponema* (ASV322, Spirochaetaceae) was significantly more abundant in NCA, where the plant taxa Marantaceae and Zingiberaceae were more abundant. Hicks et al. (2018) found the same correspondence between these gut microbial and dietary taxa in western lowland gorillas and suggested that it was due to the high fiber content of these fallback foods, which are also important for Grauer’s gorillas at low-elevation (van der Hoek, Pazo, et al., 2021).

We detect no effects of genetic relatedness or geographic distance on gut microbiome composition, despite clear group-specific microbiome patterns. Our findings thus support previous studies that show the influence of sociality on gut microbiome composition in primates (chimpanzees, Degnan et al., 2012; Moeller et al., 2016; baboons, Tung et al., 2015; colobus monkeys, Wikberg et al., 2020; black howler monkeys, Amato et al., 2017; sifakas, Perofsky et al., 2017, 2021; Rudolph et al., 2022; humans, Dill-McFarland et al., 2019) and other group-living animals (*e.g.* bighorn sheep, Couch et al., 2020). Members of the same social group travel together and experience the same environments over extended periods of time, which could synchronize their diet and also their microbiome. The gut microbiome may in addition be directly influenced by social interactions, such as grooming and coprophagy (Amato et al., 2016; Archie & Tung, 2015; Graczyk & Cranfield, 2003). However, this does not mean that host genetics are unimportant, as longitudinal studies in Amboseli baboons have shown that the primate gut microbiome is highly heritable, which cannot easily be detected in shorter-term studies (Grieneisen et al., 2021).

The plasticity of the gut microbiome supports its potential role in facilitating adaptation to different ecological conditions, which has important consequences for species evolution, dispersal and conservation. Adaptation to changes in ecological conditions as a result of climate change, range expansion, or population dispersal into novel habitats may be supported by the ability to digest diverse foods. Several studies have reported habitat-biased dispersal in mammals, including in mountain gorillas (Guschanski et al., 2008), where individual dispersal decisions appear to be driven by the availability of familiar foods. If the microbiome was implicated in restricting dietary choice, we would expect much greater conservation of dietary items across populations than what we observe here, particularly as similar food plants appear to be available in different regions. This means that gut microbiome flexibility may provide the necessary support for translocations of individuals or populations into different habitats, which is an open question in conservation management (West et al., 2019). Nevertheless, the gut microbiome may impose constraints on the diet by driving selection of foods of similar nutrient content, even if they differ taxonomically. For example, giant pandas have typical carnivore gut microbiomes despite being bamboo specialists because the nutritional value of consumed bamboo is similar to that of meat (Nie et al., 2019). Similarly, the gut microbiome of wild rhesus macaques is strongly correlated to seasonal patterns of macronutrient intake, but not food type (*e.g.*, fruit, leaves, etc.) (Cui et al., 2021). Metabolic analyses of gorilla diet, as performed for different social groups and seasons in other gorilla species (*e.g.,* Gomez et al., 2015; Rothman et al., 2008), will enable investigating whether nutritional values are conserved in different populations.

### Understanding diet and ecology of wild animals requires a combination of approaches

As every method, the metabarcoding approach to diet and microbiome faces limitations, specifically in the form of marker gene selection, reference database bias, threshold decisions, and interpretation of abundance. While Grauer’s gorillas predominantly feed on vegetative plants, they occasionally consume insects and fungi (van der Hoek, Pazo, et al., 2021; Yamagiwa et al., 1991). By choosing a chloroplast gene, *trn*L, we restricted dietary characterization in this study to plants only. For dietary analysis of species with omnivorous diets, expanding to multiple loci that are able to characterize the diversity of consumed foods would be necessary (Taberlet et al., 2018). Further, metabarcoding relies on a reference database for taxonomic identification, making it limited by the content of these databases, which may be incomplete for biodiversity-rich or extreme habitats and unstudied microbiomes (Hird, 2017; Taberlet et al., 2018). Similarly, chosen thresholds for sequence identity and relative abundance could remove genuine dietary or microbial constituents. We used conservative sequence identity and relative abundance thresholds similar to those employed in other studies of diet and gut microbiome (Deagle et al., 2019; Hibert et al., 2013; Quéméré et al., 2013; Srivathsan et al., 2016). However, this does not completely guard against removal of genuine taxa, particularly for dietary characterization, due to the small size and high variability of the *trn*L locus. Additionally, estimated abundances of the different plant and microbial taxa may not accurately reflect their abundances (Deagle et al., 2019; Gloor et al., 2017). DNA copy number can be biased by plant tissue type (*i.e.,* fruit, pith, leaves, the latter of which contain more chloroplasts; Egea et al., 2010), the copy number of the rRNA locus, relative digestibility (*i.e.,* amount of fiber), and PCR amplification success (reviewed by Deagle et al., 2019). However, other methods for dietary characterization also face biases. For example, accuracy of macroscopic fecal analysis depends on the types of tissues consumed and the extent of digestion (King & Schoenecker, 2019). Observational studies can overestimate the dietary importance of foods with longer handling times (Matthews et al., 2020) and require habituating study animals, which may make them more vulnerable to poaching and increase exposure to human-transmitted diseases (Green & Gabriel, 2020). Hence, understanding ecological and particularly dietary diversity of different animal species would benefit from a combination of approaches. Molecular methods are particularly suited for the study of unhabituated animals, in regions where tracking over a long time period is not feasible or desirable.

The use of shotgun metagenomics will ameliorate many of the limitations described above and allow for more complete interpretation by also enabling functional characterization of gut microbial communities. It would thus be feasible to test if the gut microbiome differs in functional profiles as a result of dietary differences across populations, or if functions remain conserved, suggesting that nutritional values of different diets are indeed similar. With the decrease in sequencing costs and massive growth of whole genome reference databases that become available as a result of genome sequencing initiatives (Formenti et al., 2022; Lewin et al., 2018), the use of shotgun metagenomics will increase in the coming years, fueling the application of the hologenomic framework to wild animal populations.

## Conclusions

Our results suggest that the animal gut microbiome may contribute to adaptation to new environments, while retaining a core set of potentially essential constituents. We provide evidence that this microbial plasticity is associated with dietary flexibility, and as such the gut microbiome may enable the host to exploit new resources, a precursor to local adaptation. If so, the microbiome may indirectly encourage subsequent cultural adaptation to feeding on new dietary items. We emphasize the utility of fecal sampling for minimally-invasive population monitoring of different aspects of endangered species biology, from genetics to ecology and foraging behavior. Despite its limitations, a molecular approach can reveal otherwise clandestine insights into the biology of elusive animals and is particularly powerful when combined with traditional observational methods. Our results highlight the importance of incorporating multiple axes of population differentiation into studies of endangered animals, since safeguarding ecological and genetic biodiversity is the primary objective of species conservation.

## Acknowledgements

We thank L’Institut Congolais pour la Conservation de la Nature (ICCN) and local landowners and community members for permitting us to work in the Kahuzi-Biega National Park and the Nkuba Conservation Area. We are indebted to numerous rangers and field assistants in Kahuzi-Biega National Park as well as field assistants, gorilla trackers, and local community members in Nkuba Conservation Area who, through their effort and commitment, support this project and ensure the survival of the Grauer’s gorillas. This research was supported by the Royal Physiographic Society of Lund Jan Löfqvist and Nilsson-Ehle Endowments to KG, the Erasmus Mundus Master Programme in Evolutionary Biology Consortium Scholarship to AM, and the Swedish Phytogeographical Society and the Extensus Foundation Grants to LP. The Dian Fossey Gorilla Fund’s work in DRC was supported by individual donations and grants from the Great Ape Conservation Fund of the US Fish and Wildlife Service, the Arcus Foundation, the Daniel L. Thorne Foundation, and the Turner Foundation. Sequencing was performed by the SNP&SEQ Technology Platform in Uppsala. The SNP&SEQ Technology Platform is part of the National Genomics Infrastructure Sweden and Science of Life Laboratory. The SNP&SEQ Platform is also supported by the Swedish Research Council and the Knut and Alice Wallenberg Foundation. We also acknowledge the National Bioinformatics Infrastructure for providing computational resources to this project.

## Data Accessibility & Benefit-Sharing Statement

### Data Accessibility

Sequences and associated metadata for the gut microbiome and diet generated in this project have been uploaded to the European Nucleotide Archive (ENA) under Accession no.: PRJEB49814.

### Benefit-Sharing

This research addresses a priority concern, the conservation of a critically endangered species. This was made possible by maintaining long-term collaborations with scientists in the DRC, and all collaborators are included as co-authors. Benefits of this research include the sharing of our data (above) and results with the broader scientific community as well as with conservation practitioners.

## Author Contributions

AM, RM and KG planned the study design. AM generated dietary data and performed all analyses. RM generated gut microbial data. PN, YL, MAG, and JS generated gorilla genotyping data. KN and LP provided reagents and expertise for dietary analyses. NI, AP, UN, EB, RNP, DC, and KG collected fecal samples and provided support in the field. DC, LP and KG provided project supervision. KG supervised the experiments and data analyses. AM and KG wrote the manuscript, with contribution from all authors. All authors reviewed and approved of the final manuscript.

