## Supplemental Material for "Isolated Grauer’s gorilla populations differ in diet and gut microbiome"

**Table of Contents:**


| | |
|---|---|
| Supplemental Text | This document, pg 2 |
| Supplemental Figures | This document, pg 3 - 11 |
| Supplemental Tables | Separate Excel spreadsheet |

### Supplemental Text

*Results*

#### Diet sampling depth

Sample accumulation curves suggested that additional samples would uncover only few novel *trn*L variants (**Figure S5A**), except in MNP, where only five individuals were sampled. Although deeper sequencing could have potentially uncovered novel taxa (**Figure S5B**), sequencing depth did not have a strong effect on *trn*L richness (three additional taxa per 47,000 reads, $p$ = 0.03). Furthermore, the discrepancy per sample in observed minus predicted richness did not correlate with sequencing depth ($p$ = 0.3), nor with any of the tested variables (population $p$ = 0.9; social group $p$ = 0.06; sex $p$ = 0.6; age class $p$ = 0.8; altitude $p$ = 0.4). Taken together, this suggests that our dataset is suitable to carry out comparisons across study populations.

### Supplementary Figures

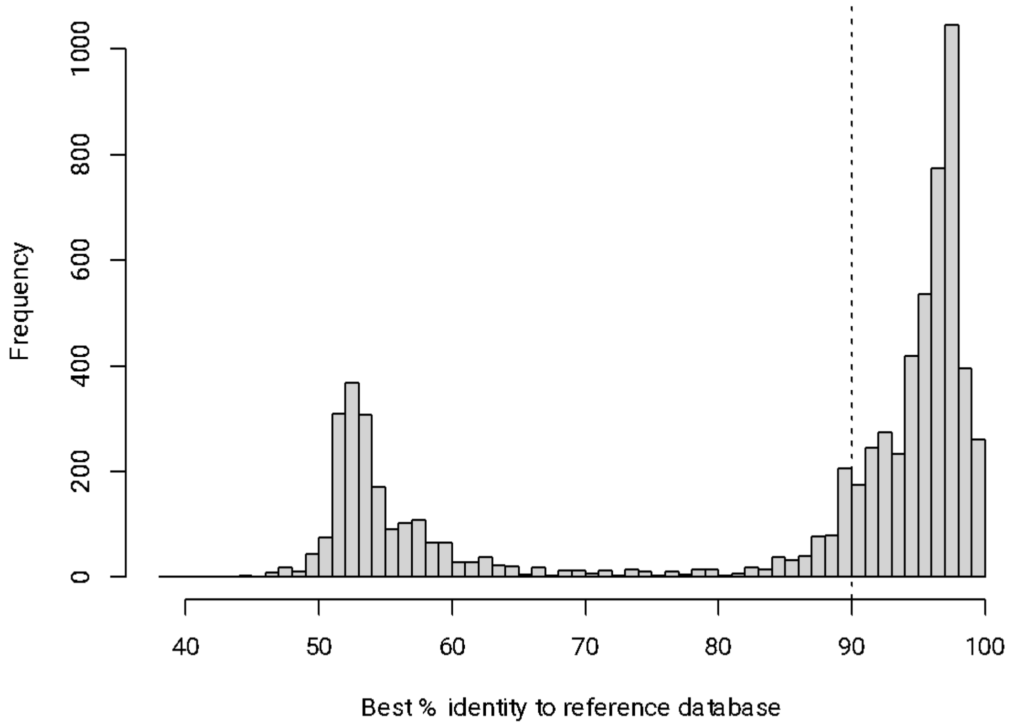


**Figure S1.** Distribution of percent identity of *trnL* reads to custom reference database. Based on this histogram and similar studies (e.g., Shehzad et al. 2016), we concluded that sequences below ~90% identity are likely chimaeras or other PCR and sequencing errors.

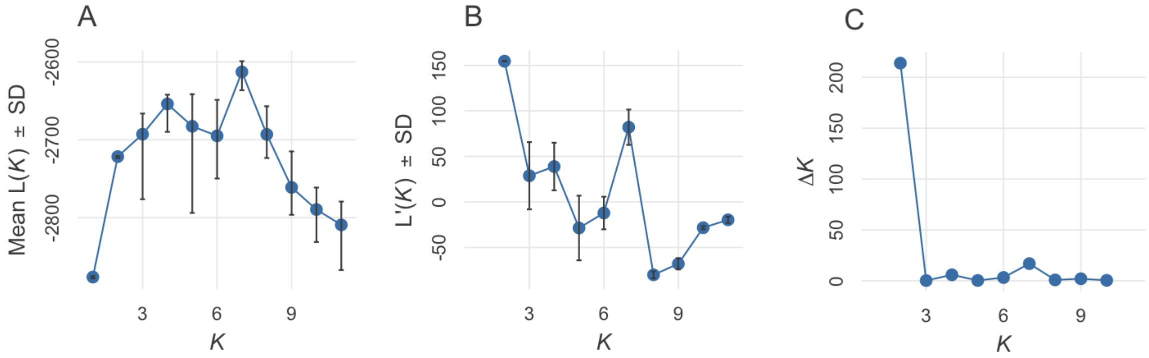


**Figure S2.** Derivative plots for STRUCTURE runs of K:1-11: (A) mean estimated log likelihood over 20 runs per K, (B) rate of change of the likelihood distribution, calculated as $L'(K) = L(K) - L(K - 1)$, and (C) Evanno's $\Delta K = m|L''(K)|/s[L(K)]$ (Evanno *et al.* 2005).

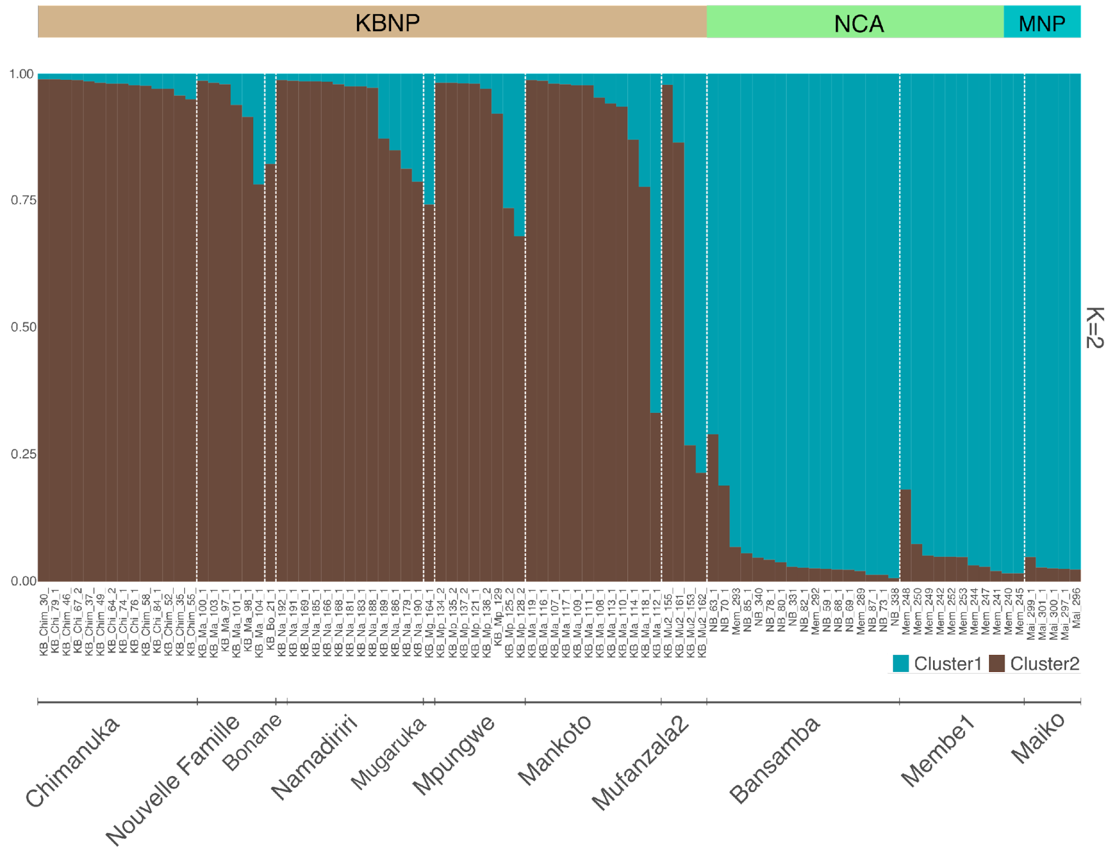


**Figure S3.** Representative run of K=2 STRUCTURE admixture plot, with each sample coloured according to inferred cluster. Plot produced using *pophelper* package in R.

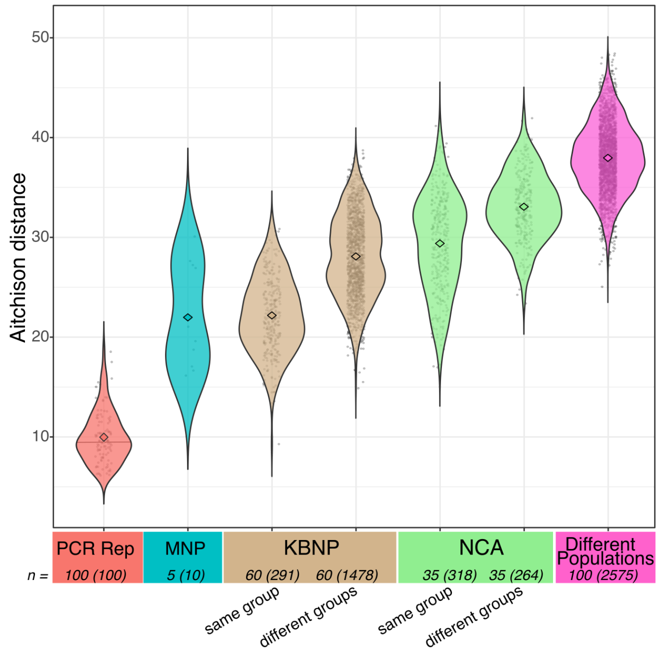


**Figure S4.** Pairwise Aitchison distances in dietary (*trnL*) composition between samples. Diamonds inside the violin plots indicate group means. *n = number of samples (number of pairwise comparisons).* PCR replicates (red) of the same sample were more similar to each other than to any other sample (Wilcoxon Rank Sum Test $p < 0.001$). Dietary profiles of individuals from the same social group were significantly more similar to each other than to dietary profiles of individuals from different social groups from the same population ($p < 0.001$). Populations showed the greatest difference in dietary profiles ($p < 0.001$).

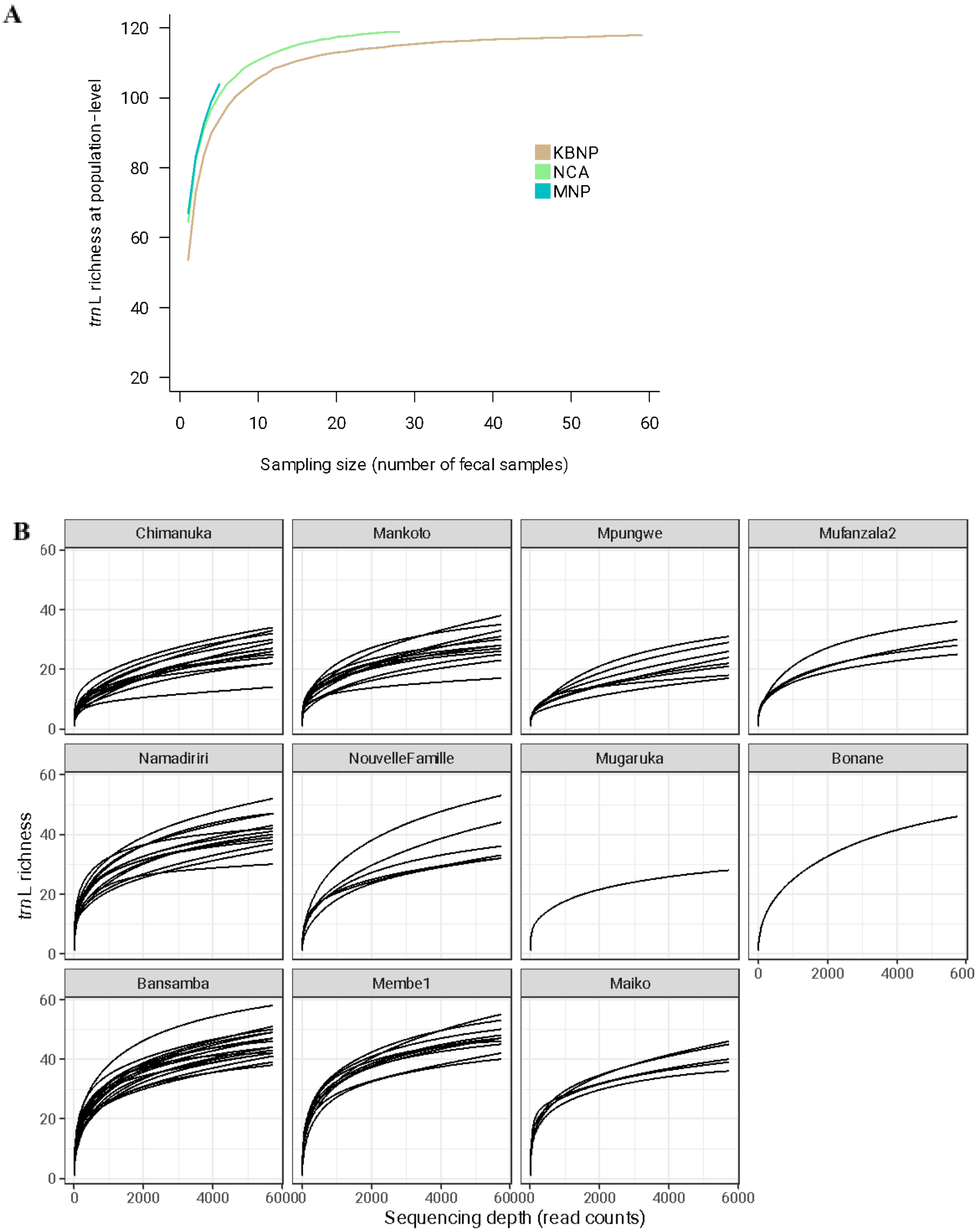


**Figure S5.** *trnL* accumulation curves (A) per faecal sample and (B) per sequencing read (equal to rarefaction curves). (A) MOTU variant accumulation curves as a function of the number of faecal samples per population ([95% confidence level]: KBNP 118 vs. 122.5 [118.50- 158.92], NCA 119 vs. 121.0 [119.18- 141.13], MNP 103 vs. 128.0 [109.72-204.72). Shaded regions represent confidence intervals over 100 random permutations using the Chao extrapolation method to estimate the number of variants (Hsieh *et al.* 2016). (B) Rarefaction curves for each social group rarefied to minimum sample sequencing depth (5743 *trnL* reads). Separate lines represent unique gorilla individuals. Not all curves reach their asymptote, suggesting that more variance may be discovered with deeper sequencing.

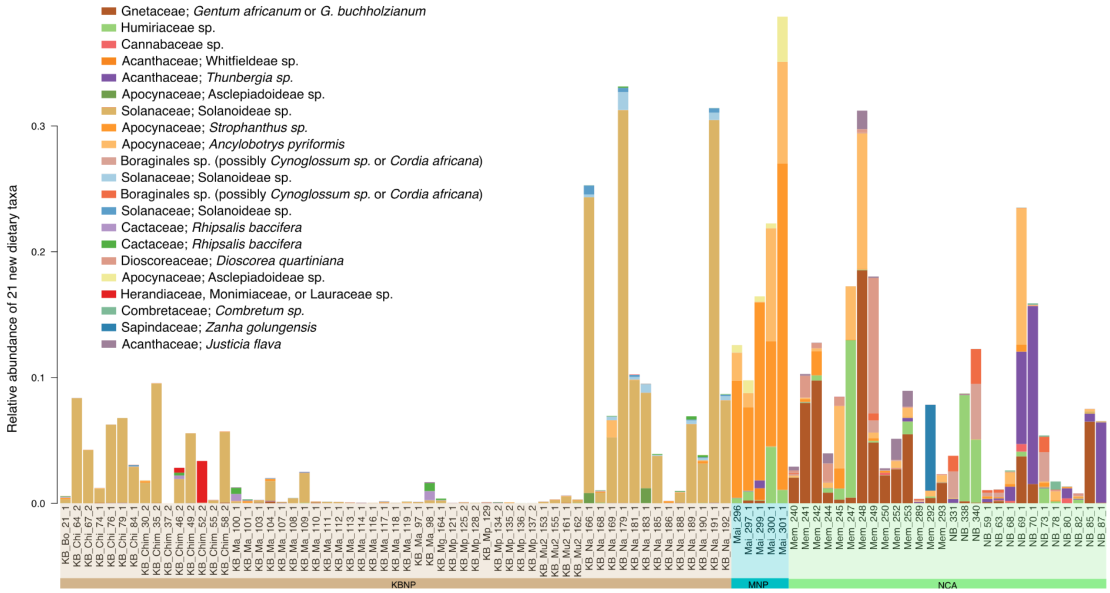


**Figure S6.** Relative abundance of 21 new dietary plants detected in Grauer's gorilla fecal samples across the three populations. While some were restricted to single samples and/or at low abundance, some taxa were abundant (*e.g.*, Solanoidea sp., *Gnetum sp.*, Whitfieldeae sp., Humiraceae sp., *Thunbergia sp.*).

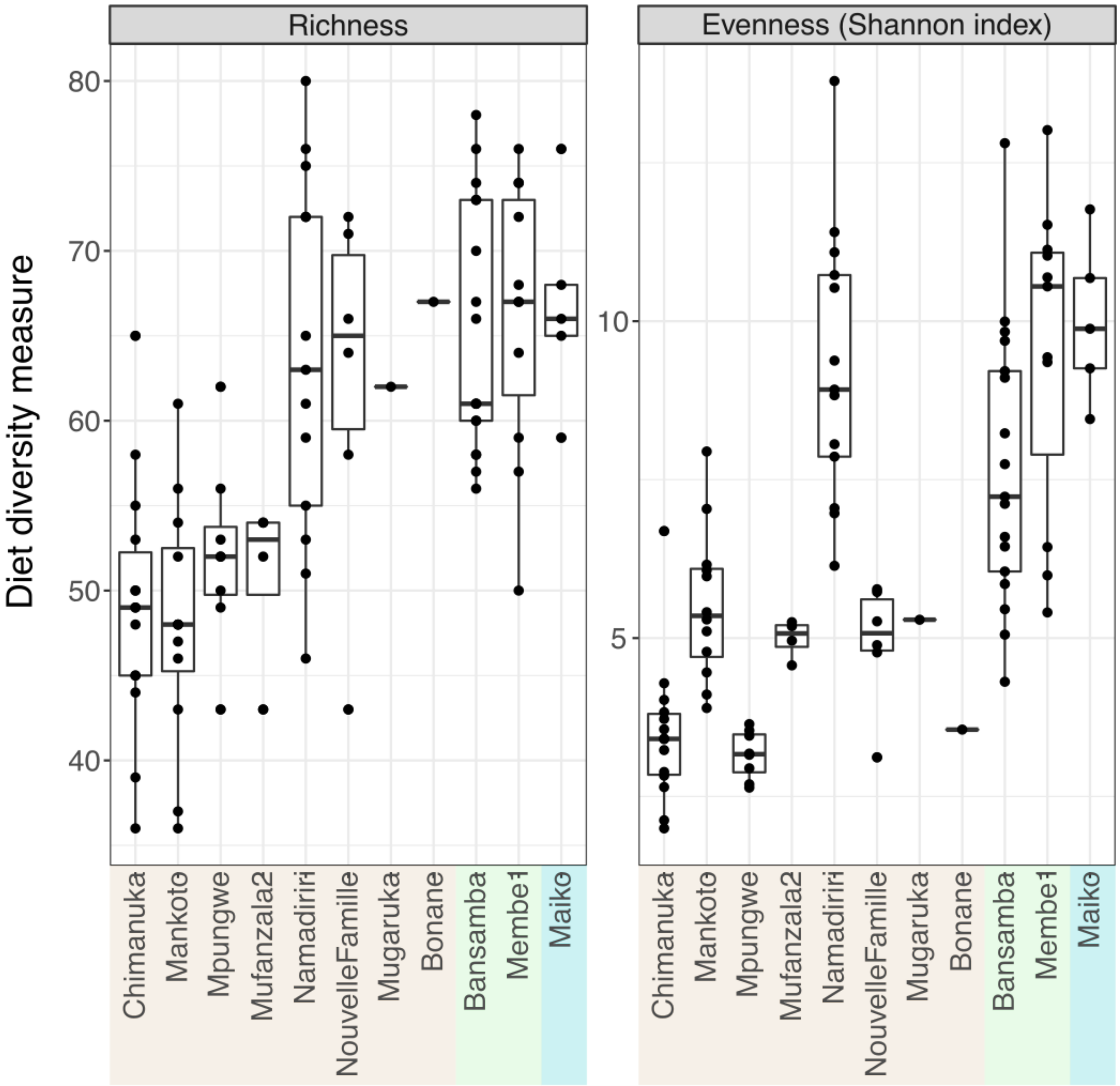


**Figure S7.** Dietary richness and evenness in gorilla social groups. Each point corresponds to an individual sample, grouped by social group or lone silverbacks (Bonane and Mugaruka). The left panel shows *trnL* taxa richness (variant or MOTU counts) per sample. The right panel shows the exponential Shannon-Wiener entropy index as a measure of dietary composition evenness. Overall, NCA samples show higher richness and evenness than KBNP samples (GLM $p < 0.001$). Pairwise statistical tests by social group, population, age class, and sex are shown in Supplemental Table 8.

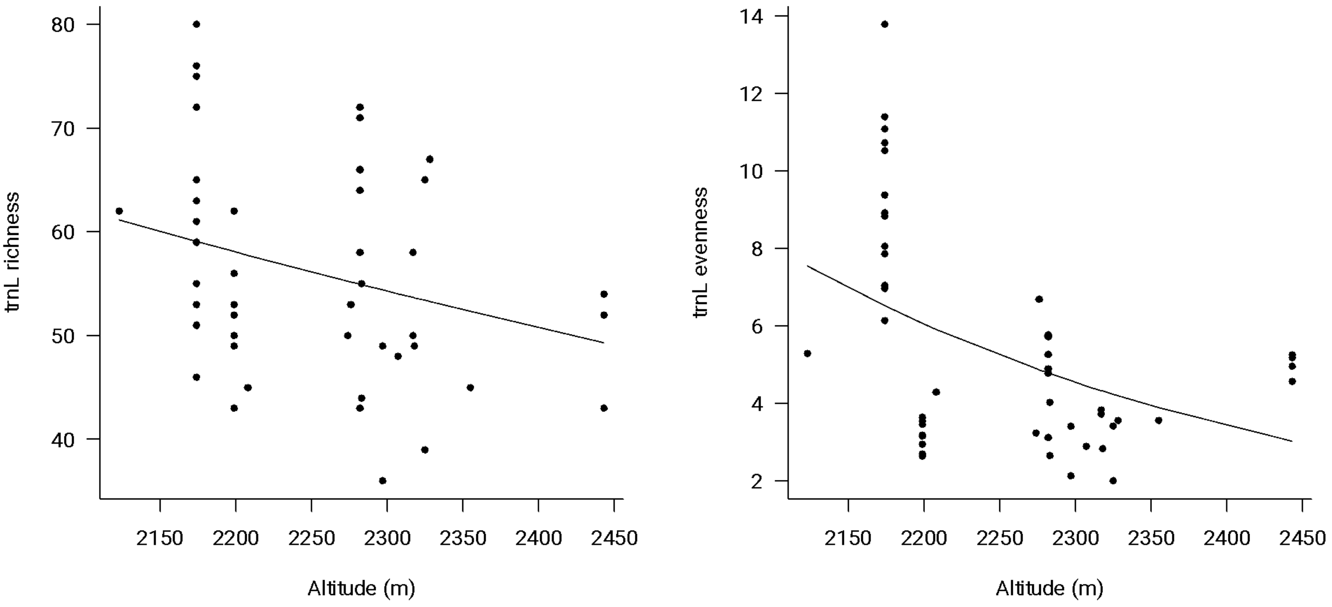


**Figure S8.** In KBNP, dietary richness (left) and evenness (Shannon true diversity index, right) declined with increasing altitude ($p < 0.001$; gamma GLM including effects of sequencing depth, sex, age class, and altitude in meters). The effect amounted to a loss of about 3.0 taxa (richness) or 1.1 effective taxon (evenness) per one standard deviation increase in altitude.

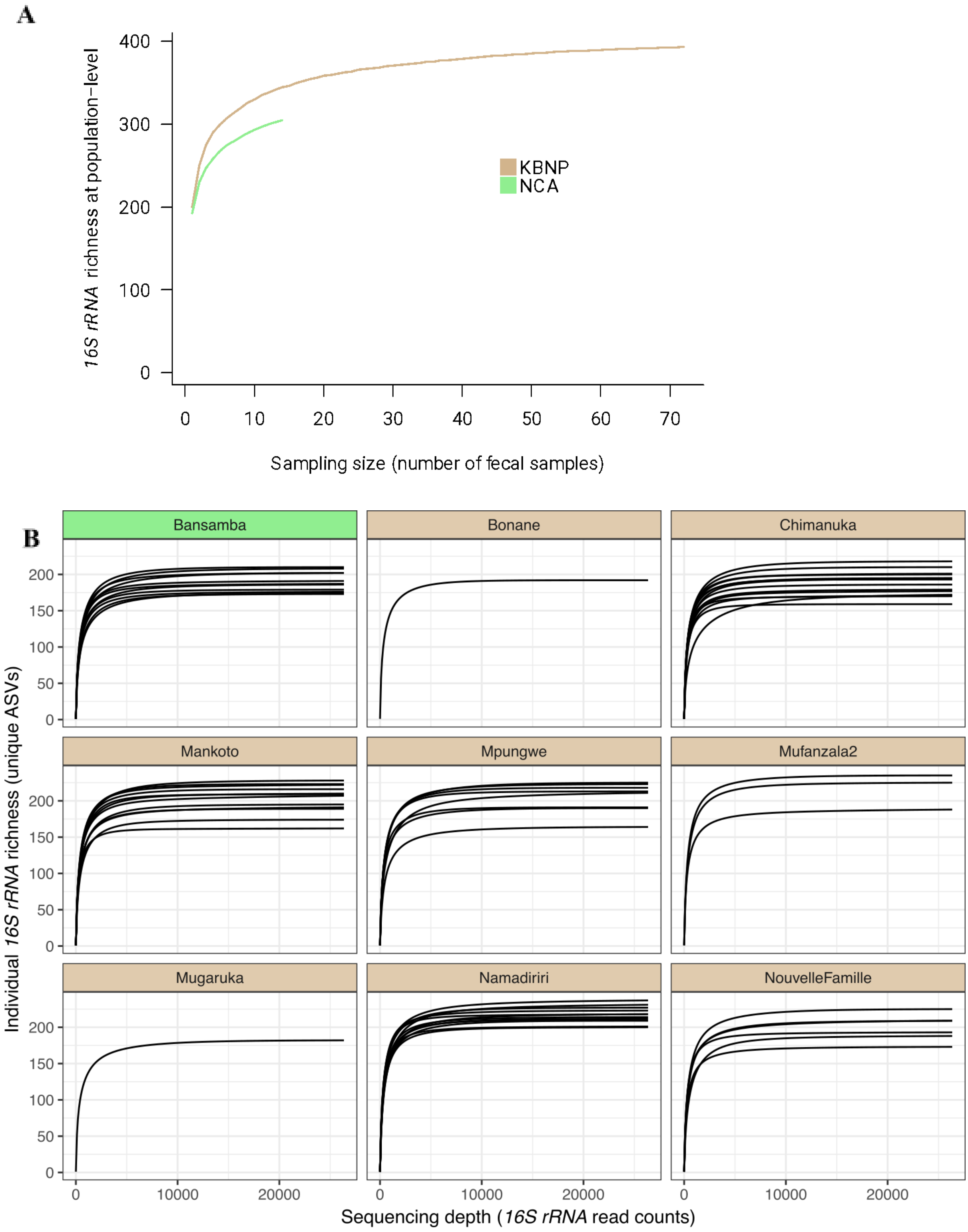


**Figure S9.** Accumulation curves for unique *16S rRNA* gene sequence variants (ASVs) at the (A) population level and (B) individual sample level based on (A) number of faecal samples collected per population and (B) number of sequence reads (rarefaction). The asymptote for (A) is estimated (Chao *et al.*, 2014) at 403.9 ASVs in KBNP (vs. 388 observed in N=57) and 347.9 ASVs in NCA (vs. 299 observed in N=11). (B) Rarefaction curves for each social group rarefied to minimum sample sequencing depth. Brown header designates social groups from KBNP, while green is the group from NCA. Sequencing depth is adequate and most samples reach the asymptote.

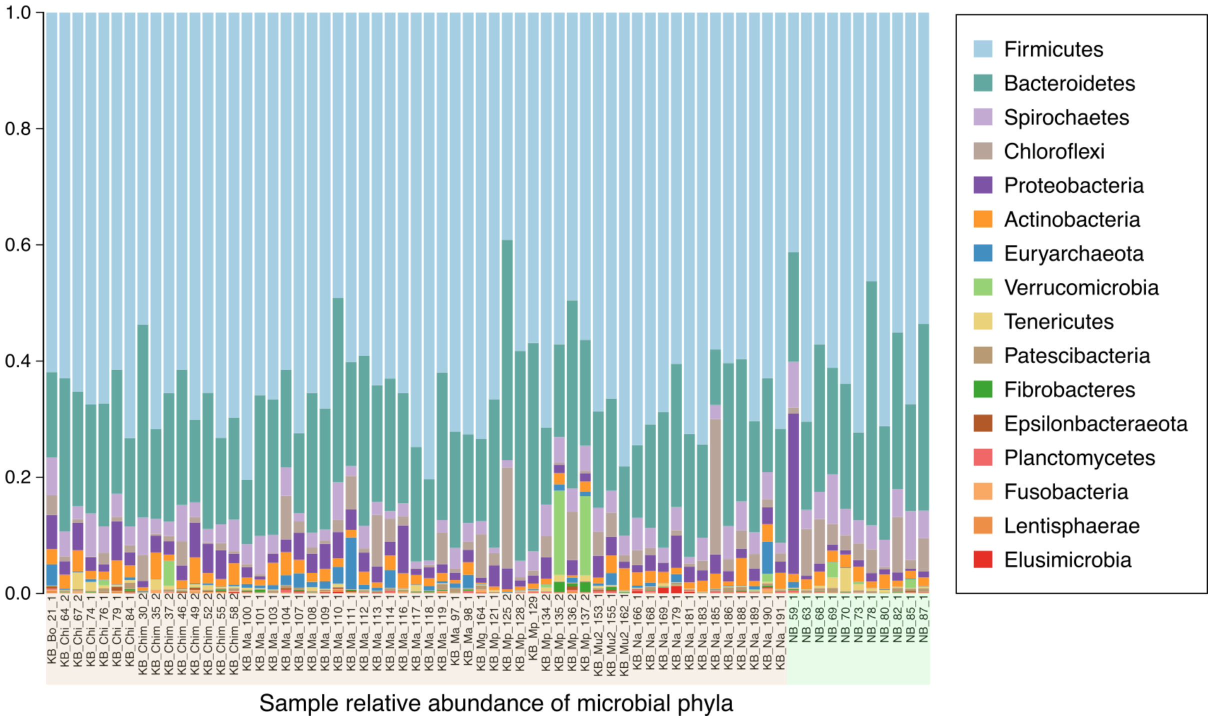


**Figure S10.** Microbial relative read abundance at the phylum level by individual sample. Sample labels are highlighted by population of origin: brown for KBNP and green for NCA.

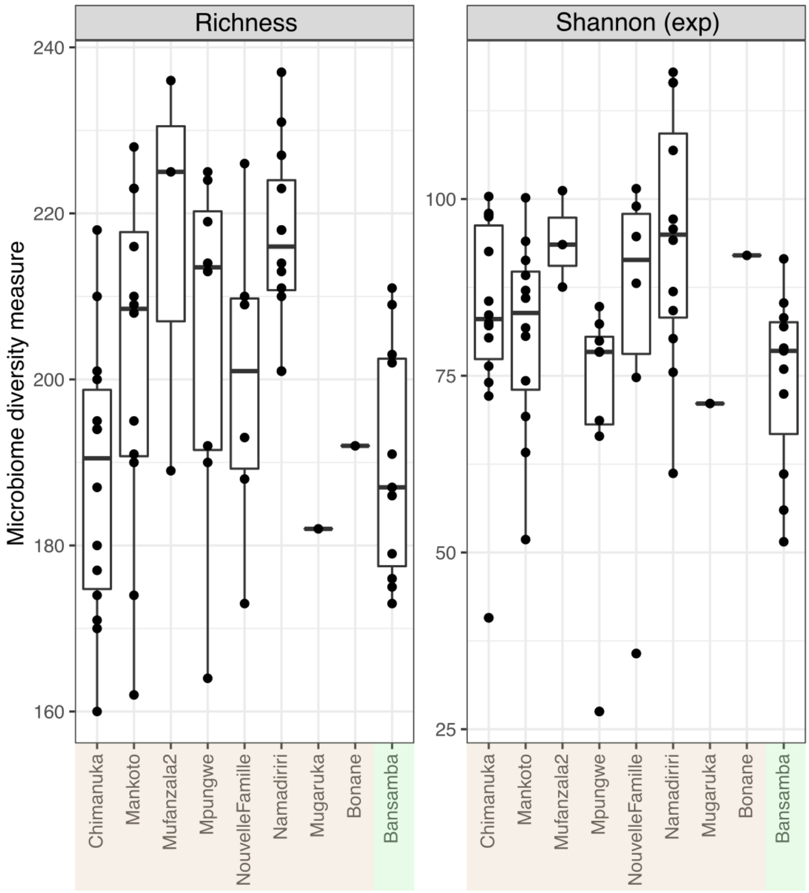


**Figure S11.** Microbiome alpha diversity metrics: number of ASVs (richness) and evenness (exponential Shannon index). Evenness was higher in KBNP than NCA ($p = 0.01$), but richness did not differ by population ($p = 0.05$).

**Supplemental Tables**

See separate Excel spreadsheet
